# Massively parallel single-cell sequencing of genetic loci in diverse microbial populations

**DOI:** 10.1101/2022.11.21.517444

**Authors:** Freeman Lan, Jason Saba, Tyler D Ross, Zhichao Zhou, Katie Krauska, Karthik Anantharaman, Robert Landick, Ophelia Venturelli

**Affiliations:** Department of Biochemistry, University of Wisconsin-Madison, Madison WI 53706; Department of Chemical & Biological Engineering, University of Wisconsin-Madison, Madison WI 53706; Department of Bacteriology, University of Wisconsin-Madison, Madison WI 53706; Microbiology Doctoral Training Program, University of Wisconsin-Madison, Madison WI 53706

## Abstract

Single cell genetic heterogeneity is ubiquitous in microbial populations and an important aspect of microbial biology. However, we lack a broadly applicable and accessible method to study this heterogeneity at the single cell level. Here, we introduce a simple, robust, and generalizable platform for quantitative and massively parallel single cell sequencing of target genetic loci in microbes using ultrahigh-throughput droplet microfluidics (Droplet Targeted Amplicon Sequencing or DoTA-seq). Using DoTA-seq, we elucidate the highly diverse single cell ON/OFF states of the phase-variable capsule synthesis operons in the prevalent human gut species *Bacteroides fragilis*. In addition, we quantify the shifts in antibiotic resistance gene abundances in different species in a 25 member human gut microbial community in response to antibiotics. By sequencing tens of thousands of single-cells derived from a human fecal sample, we identify links between plasmid replicons and the taxonomic lineages of their associated hosts. In sum, DoTA-seq is an accessible and broadly applicable tool for profiling single-cell genetic variation in microbiomes.

## INTRODUCTION

Single-cell heterogeneity is ubiquitous in nature and single-cell sequencing is a powerful tool for understanding the biology of systems composed of heterogenous cells. In bacteria, single-cell genetic heterogeneity plays key roles in evolution, antimicrobial resistance^1^, host-colonization^2^, and pathogenesis^3^. Mechanisms of genomic variation such as phase variation, gene deletion, gene duplication, and horizontal gene transfer, are frequently observed in bacteria. However, it has been difficult to study this variability without robust pipelines for single-cell sequencing. Additionally, most microbes live in multi-species microbial communities, where studying them at the single-cell level is important for understanding the functions of each species. Classically, single-cell genetic heterogeneities are observed through colony plating where single colonies represent populations usually derived from single cells^4^. However, colony plating only detects culturable microbes and thus fails to represent unculturable taxa, which can play key roles in microbial community functions^5,6^. In addition, the lack of scalability limits their use to low richness (i.e. number of species) communities, small numbers of samples, or both.

Despite the demand for them, methods for single-cell genetic sequencing of microbes are not yet widely accessible^7–9^. This deficit is due in part to the significant challenges associated with sequencing single microbial cells. The diverse makeup of their cell membranes, cell walls, and other protective features require different lysis or permeabilization conditions specific to the individual microbial species. These challenges result in single-cell sequencing protocols that require complex workflows and/or are not generalizable to diverse microbial systems^7–9^.

More recently, novel strategies have been developed based on next generation DNA sequencing to study genotypic heterogeneity in microbial communities at a finer resolution. Metagenomic sequencing in combination with techniques used in chromosome conformation mapping of eukaryotic cells (Hi-C based methods)^10,11^ have been elegantly adapted to physically link proximal genes and then elucidate gene-taxa and taxa-taxa associations within microbial communities. This method has the advantage of being untargeted, enabling a discovery-driven approach to revealing gene-taxa links within microbial communities. However, this method requires complex data analysis workflows.

On the other hand, emulsion PCR based methods, where multiple PCR amplicons from single cells are fused by PCR into a single amplicon for sequencing^12^, have been used to link genes of interest within cells in microbial populations^13,14^. However, a major limitation of these methods is that the linked amplicon is only generated when all target sequences of interest are present in the cell and correctly amplified. These methods fail to provide information about cells that are missing one or more target sequences. Furthermore, the limitations in next-generation sequencing and PCR efficiency imposes a limit on the ability to amplify and sequence long amplicons composed of many target sequences. Therefore, the limited number and length of target sequences restrict the generalizability of the method. Finally, the integration of cell lysis and PCR into a one-pot reaction severely limits the range of lysis conditions needed for efficient extraction of DNA from diverse bacterial species due to potential PCR inhibition by the lysis reagents.

Overcoming the limitations of the previous approaches, microfluidics-based barcoded single-cell whole genome sequencing holds promise as a generalizable and quantitative method for profiling single-cell genotypic heterogeneity^9,15^. However, the complex microfluidic workflows (droplet reinjection and droplet merging) and high sequencing costs per cell make this method most suited for large-scale genome sequencing projects backed by substantial resources. Thus, widespread deployment of these methods to scientific community at large is limited. Thus, despite these recent advances, it remains challenging for most labs to study microbial single-cell genetic heterogeneity. As such, there is a pressing need for a widely accessible, generalizable, and quantitative method for profiling genetic heterogeneity within microbial populations at the single-cell level.

Here, we report a robust, and generalizable droplet microfluidics workflow for quantitative single-cell targeted genetic profiling of microbes (Droplet Targeted Amplicon Sequencing or DoTA-seq). DoTA-seq uses a microfluidic droplet-maker to encapsulate single microbial cells into millions of picoliter-sized hydrogels. This method separates the steps of cell lysis and PCR which in turn allows multi-step cell lysis procedures without multiple microfluidics steps. Furthermore, it uses multiplexed PCR to attach unique DNA barcodes to the targeted genetic loci of each cell, providing flexibility in the number and type of loci targeted for sequencing. Notably, DoTA-seq leverages the advantages of ultrahigh-throughput droplet microfluidics, allowing quantitative single-cell sequencing of microbes without requiring complex microfluidic processes such as droplet re-injection and droplet merging. This workflow relies only on microfluidic droplet makers and readily available reagents. Hence, it has the potential to be adopted widely by the biology research community.

To demonstrate the broad utility of DoTA-seq, we simultaneously and quantitatively profile genetically distinct subpopulations generated by phase variation in the prevalent human gut symbiont *Bacteroides fragilis* (*B. fragilis*). In addition, we tracked the taxonomically resolved frequency shifts of 12 antibiotic resistance genes within a 25-member human gut community exposed to increasing concentration of antibiotics. These data reveal temporal changes in the presence of resistance genes at the single-cell level within members of this community. Lastly, use DoTA-seq to profile the host-range of mobile genetic elements in a human fecal microbial community by sequencing up to >37,000 cells in a single condition. The highlighted applications are only possible using DoTA-seq and represent just a few of many possible future applications of this workflow. In sum, DoTA-seq is a powerful and generalizable method for studying microbial genomes at a single-cell resolution.

## RESULTS

### The DoTA-seq workflow

The power of droplet microfluidics to isolate and carry out reactions has been applied to single-cell sequencing^9,15–17^. However, one significant draw-back of droplet microfluidics is the complexity of multi-step workflows. In particular, these methods require a combination of droplet reinjection, splitting, and merger steps which have high failure rates and are difficult to implement without extensive droplet microfluidics expertise^9,18,19^. As a result, compromises in efficiency are typically made to achieve cell lysis and molecular reactions simultaneously in a one-pot reaction^14^.

For many microbial communities, the diversity of bacterial cell walls precludes a one-pot reaction that efficiently lyses all species, while allowing for the desired molecular reactions to occur. By encapsulating microbial cells in hydrogels matrices^9,13,20^, we can use detergents and lytic enzymes to achieve multi-step lysis while maintaining intact genomic DNA. In DoTA-seq, we encapsulate single cells in reversibly crosslinked polyacrylamide hydrogels **(Fig. 1a)**. This type of hydrogel is heat resistant, enabling a wide repertoire of potential lysis conditions to achieve efficient lysis of diverse microbes. The pore size of this polymer matrix is small, retaining small genetic elements such as extrachromosomal plasmids.

**Figure 1.**
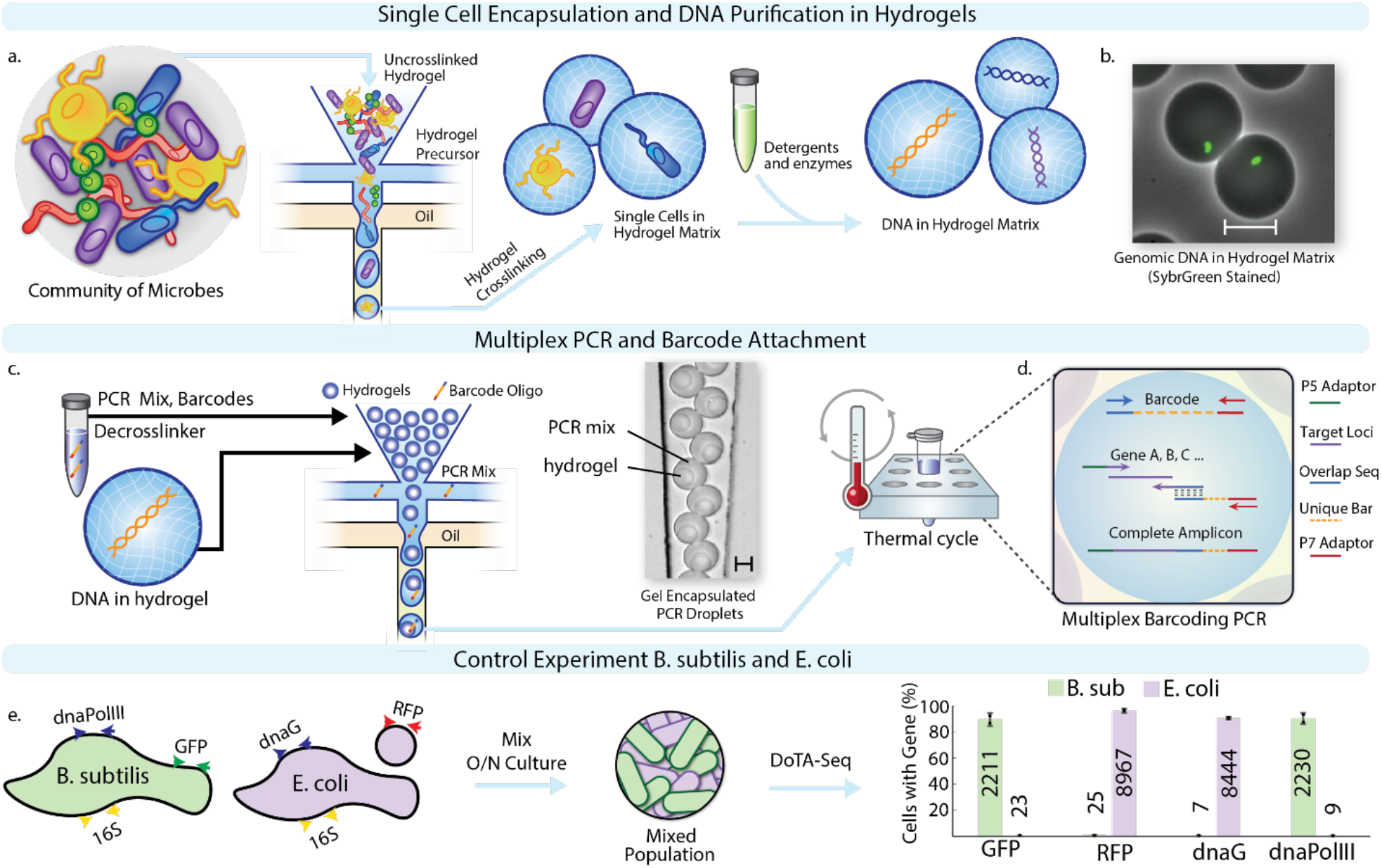
Droplet Targeted Amplicon Sequencing (DoTA-seq) accurately profiles single-cell genetic loci in gram-negative and positive bacteria. **(a)** Overview of the DoTA-seq workflow. Using a microfluidic droplet maker, microbes are encapsulated in a hydrogel matrix according to a Poisson distribution such that most hydrogels contain zero or one cell. Chemical, enzymatic, and/or heat treatments are used lyse the cell within the hydrogel, leaving the genomic and plasmid DNA trapped inside the hydrogel matrix. **(b)** An overlayed fluorescent and brightfield image following lysis of *B. subtilis* cells inside hydrogels. SYBR green staining reveals the trapped genomic DNA inside the hydrogels. Scale bar represents 20μm. For images of the complete lysis workflow, see **Supplementary Fig. 4. (c)** These hydrogels are re-encapsulated with PCR mix containing PCR reagents and primers targeting specific loci, hydrogel de-crosslinker, and unique barcode oligos at a limiting dilution. Microscopy image shows the re-encapsulated gels at the outlet of the microfluidic device. Scale bar represents 20 μm. **(d)** Schematic of the barcoding multiplex PCR inside droplets. The target loci and the DNA barcodes are simultaneously amplified during PCR. Complementary regions between the forward primer of the barcode and reverse primer of the target locus results in an overlap-extension generating a spliced product compatible with Illumina sequencing. **(e)** Monocultures of *B. subtilis* and *E. coli* are mixed then sequenced using DoTA-seq, targeting a designed set of chromosomal and plasmid loci. Grouped bar plot (both species for each gene side-by-side) shows the fraction of cells with the respective gene detected in *B. subtilis* (green bars, ∼2,000 cells) or *E. coli* (red bars, ∼10,000) cells classified by 16S rRNA sequencing. Numbers on bars represent average number of cells detected. Dots represent independent values and error bars represent 1 s.d. from the mean of technical replicates (n=2). O/N denotes overnight cultures.

To enable reliable and robust amplification of target sequences from single cells, we first encapsulate bacterial cells at limiting dilution into water-in-oil droplets using a microfluidic droplet maker such that most droplets contain one or zero cell according to a Poisson distribution (**Fig. 1a, Supplementary Note 1**). At limiting dilution, approximately one in ten droplets will contain a cell, but the microfluidic droplet maker operates at ∼10 kHz frequencies, encapsulating hundreds of thousands of single cells within a few minutes. The droplets contain acrylamide monomers and crosslinkers, which are polymerized and reversibly crosslinked into a polyacrylamide matrix, trapping individual cells within the matrix. The polyacrylamide gel matrix contains pores on the scale of 10-100nm^21^, allowing detergents and enzymes to diffuse through, lysing the cells and removing cellular material while entrapping genomic and plasmid DNA (**Fig. 1b**).

Next, we re-encapsulate the gels containing the lysed cells with PCR reagents and random DNA barcodes at a limiting dilution such that most droplets contain one gel and one or zero barcodes according to a Poisson distribution (**Fig. 1c**). Typically, we use a Poisson loading ratio of ∼0.1, which means approximately one in ten droplets contain one barcode, and approximately 0.5% of droplets contain more than one barcode. A droplet containing more than one barcode can result in cells that are counted more than once, but since these events are low frequency and occur randomly, they do not substantially impact the sequencing results. Correspondingly, throughput could be augmented with some loss in quantitative resolution simply by increasing the Poisson loading ratio of the barcodes (i.e. fewer droplets contain zero barcodes, but a higher fraction of droplets contain multiple barcodes).

In droplets that contain a lysed cell and a unique barcode, multiplex PCR simultaneously amplifies both the target genetic loci and the barcode and splices them together into an amplicon library for sequencing on Illumina platforms (**Fig. 1d**). In droplets without lysed cells or barcode, complete amplicons are not generated, and are thus not sequenced. In sequencing analysis, a matching barcode sequence signifies amplicons that derive from the same droplet and thus the same cell. Rare events from Poisson loading, irregularities in droplet making, and coalescence of droplets during PCR can result in multiple cells associated with the same barcode sequence. These events (i.e. barcodes) can often be filtered out based on unique signatures during data analysis **(Supplementary Note 2)**.

The microfluidic droplet makers used in DoTA-seq are simple in design and easy to use and fabricate. Similar devices can be purchased from commercial sources (**Supplementary Note 3**). All reagents for this workflow are widely available off-the-shelf. In our lab, this workflow is regularly performed for 5 samples at a time for ∼10,000 cells per sample pooled into one Miseq run, which required <8 hours of hands-on time starting from initial sample to collecting sequence data.

### DoTA-seq efficiently captures multiple genetic loci in gram-positive and gram-negative bacteria

To evaluate the ability of DoTA-seq to capture target loci in gram-negative and gram-positive bacteria, we used a defined mixture of *Escherichia coli* MG1655 and *Bacillus subtilis PY79. E. coli* contained a 4kb plasmid harboring red fluorescent protein (*RFP*) and a pSC101* origin and *B. subtilis* harbored a chromosomally integrated green fluorescent protein (*GFP*)^22^. We mixed equal volumes of both species and performed DoTA-seq on the mixture. To determine whether we could measure multiple loci on the chromosomes and plasmid, we targeted *RFP* on the plasmid, *GFP* on the chromosome, and the endogenous genes *dnaG* and *dnaPolIII* for *E. coli* and *B. subtilis*, respectively. In addition, we targeted the 16S rRNA V3-V5 region for both species (**Fig. 1e**).

We sequenced a total of ∼10,000 cells per run (2 technical replicates per sample). To determine whether the targeted genes were faithfully linked to the correct species, we first aggregated all reads with the same barcode to represent a given cell. Since the 16S rRNA gene sequence is widely used as an indicator for taxonomy, we used the 16S sequence to classify the species identity of each cell. We then aggregated all barcodes associated with a single species and counted the fraction of cells that contained reads mapping to each target gene. For both *E. coli* and *B. subtilis* cells, we detected the expected genes in ∼90% of the sequenced cells, showing that DoTA-seq can efficiently capture target genes within single cells (**Fig. 1e, Supplementary Fig. 1**). At the same time, false positives (genes not belonging to the assigned species) were detected in ∼0.2% of cells for the chromosomal genes (*dnaG, dnaPolIII*, and *GFP*) and ∼1% for the *RFP* on the plasmid. The ∼4kb plasmid is orders of magnitude smaller in length than genomic DNA and is also present in multiple copies per cell. Therefore, the plasmid has a higher chance of diffusing out of its cognate hydrogel into neighboring gels during the washing, lysis, and storage steps. As such, sequences on plasmids will have higher false positives rates than chromosomal sequences. This higher false positive rate imposes a higher limit of detection for plasmid.

To further assess the ability of DoTA-seq to analyze mixtures of diverse bacteria, we used DoTA-seq to sequence a commercially available microbial community standard (ZymoBIOMICs community standard). This is a standardized community containing a wide range of characterized microbial species at pre-determined relative abundances that was characterized using metagenomic shotgun sequencing. Based on available genome sequences of the species in the sample, we designed primers targeting one essential gene for each of the species observed as well as the 16S rRNA gene. Due to the sample being preserved in a proprietary inactivating buffer (DNA/RNA shield) that lysed gram-negative bacteria, we could only reliably detect the gram-positive species in this sample with DoTA-seq **(Supplementary Fig. 2)**. DoTA-seq was successfully applied to fresh or cryo-preserved gram-negative and gram-positive bacteria. Thus, the underrepresentation of gram-negative bacteria is a limitation of the sample preservation method.

Using DoTA-seq, we detected all gram-positive species at comparable relative abundances to those obtained by metagenomic shotgun sequencing with the exception of *S. aureus* and *L. fermentum* (**Supplementary Fig. 3a**). In addition, the targeted essential genes displayed high prevalence (**Supplementary Fig. 3b**). Since our results did not involve optimization of primer proportions, future optimization of primer concentrations/proportions **(Supplementary note 5)** is expected to further improve the gene detection rate. To evaluate the lysis efficiency, we pre-stained the cells with a dye that binds to cell walls and membrane components (Cellbrite Fix 555) and SYBRgreen (DNA stain) (**Supplementary Figure 4a**). Cellbrite (Cell) and SYBRgreen (DNA) staining showed that the fraction of unlysed cells (Cellbrite positive and SYBR positive cells, comprising ∼3.9% of droplets) was substantially reduced following lysis to 0.1% (**Supplementary Fig. 4c**). This indicates efficient cell lysis of diverse bacteria while preserving encapsulated DNA (∼3.9% of droplets were SYBR positive before and after lysis) (**Supplementary Figure 4b**). Therefore, DoTA-seq can be used to sequence diverse bacterial species and communities.

### DoTA-seq reveals diverse bacterial subpopulations generated through combinatorial phase-variation

Phase variation is a biological process to generate genetic and phenotypic heterogeneity across a microbial population by modifying gene expression through heritable and reversible genetic changes^23^. By generating heterogeneity across the population, phase variation can serve as a bet-hedging strategy in uncertain environments to promote population fitness^24^. For example, phase variation contributes to the regulation of genes involved with virulence and host-colonization^2,25,26^. Observing the subpopulations generated through phase variation at multiple loci (i.e. combinatorial phase variation) could enable a deeper understanding of the underlying mechanisms behind diversification and the impact of population diversification on fitness and functions. For example, capsular polysaccharide (CPS) are polysaccharide shells regulated by phase variation that surround bacterial cells and offer protection from antimicrobials and the host immune system^2^.

To demonstrate DoTA-seq’s ability to observe these subpopulations, we performed single-cell sequencing of the invertible loci of the capsular polysaccharide (CPS) within single cells of the prevalent human gut species *Bacteroides fragilis. B. fragilis* NCTC9343 contains 8 CPS operons, each of which can synthesize a different CPS^27^. The promoters of 7 operons are regulated by an endogenous recombinase (*mpi*) which toggles the promoter between the ON and OFF state (**Fig. 2a**). Thus, in any *B. fragilis* population, there are 128 possible CPS operon promoter states, where different combinations of promoter ON and OFF states could yield unique CPS phenotypes. Thus far, the *B. fragilis* CPS promoter orientations have been examined independently for each promoter at a population-level^27,28^. Therefore, we lack an understanding of the extent of combinatorial promoter variation across a bacterial population.

**Figure 2.**
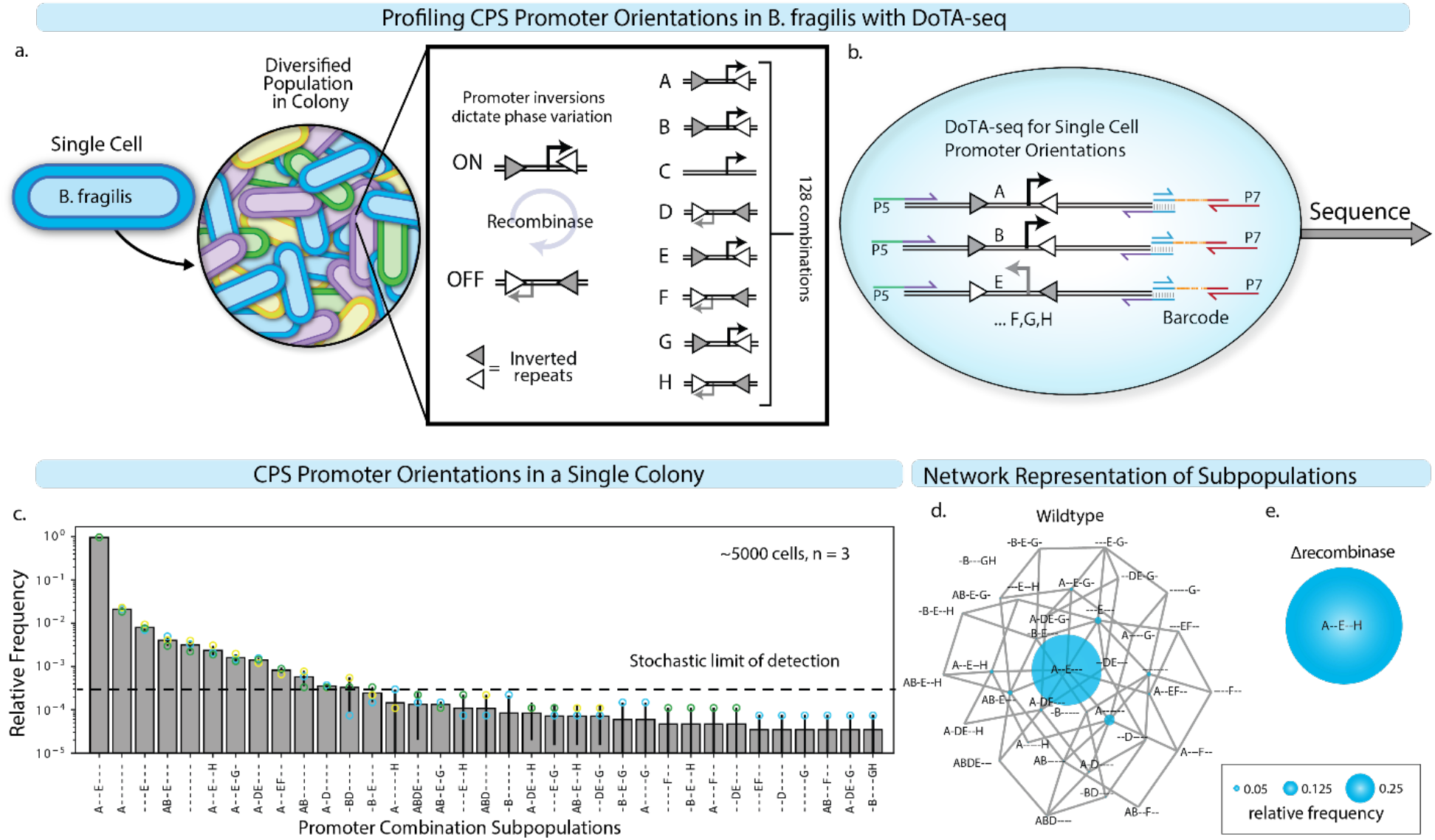
DoTA-seq elucidates diverse genetic sub-populations in *B. fragilis* generated via phase-variation of capsular polysaccharide (CPS) operons. **(a)** Schematic of the *B. fragilis* CPS operons. Phase variation in *B. fragilis* generates diverse populations starting from a single cell. There are a total of 8 CPS operons referred to as A-H, 7 of which contain promoters flanked by inverted repeats (triangles). These promoters switch ON and OFF through recombination at the inverted repeats driven by an endogenous recombinase (*mpi*). **(b)** Elucidating CPS promoter ON/OFF states using DoTA-seq with primers designed to flank all 7 invertible promoters. Primers are represented by half-headed arrows. Gray vertical lines represent region of complementarity between amplicons and barcodes. P5 and P7 represent the Illumina sequencing adaptor sequences. **(c)** Bar plot of the relative frequencies of unique CPS promoter states in a single *B. fragilis* colony. Promoter states are represented by a 7-letter code, where the letters (A-H) denote that a given promoter is turned ON, and “-” denote the given promoter is turned OFF. Data points represent technical replicates. Error bars represent 1 s.d. from the mean of technical replicates (n=3). Bar height represents the mean. Since a subset of combinatorial promoter states were rare in the population and not observed in all technical replicates, we computed the stochastic limit of detection, This detection limit is determined by the number of cells sequenced, where upon random sampling of 5000 cells, the subpopulation is expected to be detected at least 80% of the time (**Supplementary Note 4**). **(d)** An undirected graph network representation of the CPS promoter state subpopulations in (c). Nodes represent CPS promoter combinatorial states where diameter is proportional to relative frequency, and edges connect nodes that are one promoter flip away from each other. **(e)** Network representation of the measured combinatorial promoter states where the recombinase (*mpi*) responsible for promoter inversions was deleted. In this strain, the entire population is locked in a single state (A--E-H). The diameter of the node and edges are the same as (d).

To elucidate the ON/OFF states of each promoter driving a given CPS operon in each cell with DoTA-seq, we targeted each phase variable promoter with primers flanking the invertible regions (**Fig. 2b**). Since the ON/OFF orientation for each promoter produces a unique DoTA-seq amplicon, we can deduce the combinatorial promoter ON/OFF states for each cell by matching the amplicon sequence to the expected sequences for each promoter orientation. To observe the diversity of combinatorial promoter states, we quantitatively profiled the CPS promoter states in a *B. fragilis* colony grown for 48 hours on an agar plate. Since a colony is derived from a single cell with a unique promoter orientation combinatorial state, additional observed states should be the result of diversification within the 48-hour timeframe.

Since the population was composed entirely of gram-negative bacteria that did not require multi-step lysis procedures, we simplified the procedure by skipping the lysis step and directly encapsulating single cells with PCR mix (see Methods: DoTA-seq of *B. fragilis* CPS). By sequencing the colony in three technical triplicates (∼5000 cells for each replicate), we observed 35 different promoter states, which accounts for ∼25% of all possible promoter states (**Fig. 2c**). We observed low technical variation for the subpopulations that were above the stochastic limit of detection (which can be lowered by sequencing more cells). Pooling the cells from all three technical replicates (∼15,000 cells), we generated an undirected network representation of the subpopulations and their connection to other subpopulations observed in the colony (**Fig. 2d**). The largest subpopulation (promoters A and E turned ON, denoted as A--E--- where - represents B,C,F,G, or H promoters in the OFF states) constituted >90% of cells measured in the colony. Across all single cells, up to 4 promoters were observed to be simultaneously turned ON, and some cells had all invertible promoters turned OFF. Most subpopulations were within three promoter inversion steps of the largest subpopulation. In addition, every observed subpopulation was connected to at least one other subpopulation in the network except the -B---GH combinatorial promoter state, indicating that nearly all observed subpopulations were one promoter flip away from at least one other observed subpopulation.

These results are consistent with the diversification of promoter states during colony growth from an initial A—E—state, yielding new subpopulations in closely related alternative states. Based on this proposed mechanism, most of the population remained in the A--E--- state, suggesting that promoter inversion rates were slower than growth rates. In a *B. fragilis* strain deleted for the *mpi* recombinase gene^29^, all sequenced cells (∼13,000 cells) had identical promoter states (**Fig. 2e**). These results demonstrate that DoTA-seq can quantitatively resolve with high precision the single-cell genetic heterogeneity in microbial populations, making it a powerful tool for investigating phase variation. Notably, with DoTA-seq, these measurements did not require genetic modifications to the strain of interest (e.g. fluorescent promoter fusions).

### DoTA-seq reveals gene dynamics within complex microbial communities

Bacterial genomes are highly plastic, able to lose and gain genes in response to changing environmental stresses^30,31^. These gene dynamics are critical to the emergence of antimicrobial resistant pathogens^32^. Despite their enormous impacts on global health, little is known about the temporal trends in mobile genes such as antibiotic resistance genes (ARGs) and their transfer among hosts within microbial communities. This limited understanding stems from a dearth of methods for quantitatively tracking the taxonomic associations of these mobile genes (i.e. in which community members reside). DoTA-seq directly and quantitatively measures genes of interest within single cells and provides a powerful way to observe the dynamics of these genes within microbial communities.

To demonstrate DoTA-seq’s ability to track ARGs within complex microbial communities, we generated a synthetic microbial community composed of 25 prevalent members of human gut microbiota that have been extensively characterized^33^ (**Supplementary Table 1**). We identified 12 antibiotic resistance genes (ARGs) within the published genomes of these isolates and designed DoTA-seq primers targeting each ARG and each species’ 16S rRNA for taxonomic identification. Each species was grown individually, then mixed (according to approximately equal OD600, but not necessarily equal cell numbers due to variation in the OD600 to cell number relationship among species^34^) to generate the synthetic community composed of the 25 species. We performed DoTA-seq, sequencing ∼3,000 to ∼10,000 cells per sample, on this community to elucidate the ARG-species associations.

DoTA-seq revealed the expected ARG-species associations based on published genome sequences **(Fig. 3 a,b)**. One exception was the rifampicin resistant allele of the *rpoB* gene in *Bifidobacterium longum* (BL). This gene was not observed in the published BL genome sequence, but it was detected in BL single cells by DoTA-seq. However, we confirmed this gene in purified genomic DNA by PCR, suggesting that the published genome sequence was incomplete (**Supplementary Fig. 5**). Most ARGs were observed in >70% of single cells (prevalence of >70%) of a given species (*tetO* in *Anaerostipes caccae* AC, *Dorea longicatena* DL or *cepA* in *Bacteroides fragilis* BF, for example). However, some genes were observed at <50% (*tetO* in CC, DF for example). Overall, the variation in prevalence between technical replicates was small and negatively correlated (Spearman Rho = –0.57, P < 6.5e–10) to the number of sequenced cells for each species (**Fig. 3b, Supplementary Fig. 5**). This implies that increasing the total number of sequenced cells can reduce some of the technical variability observed in DoTA-seq.

**Figure 3.**
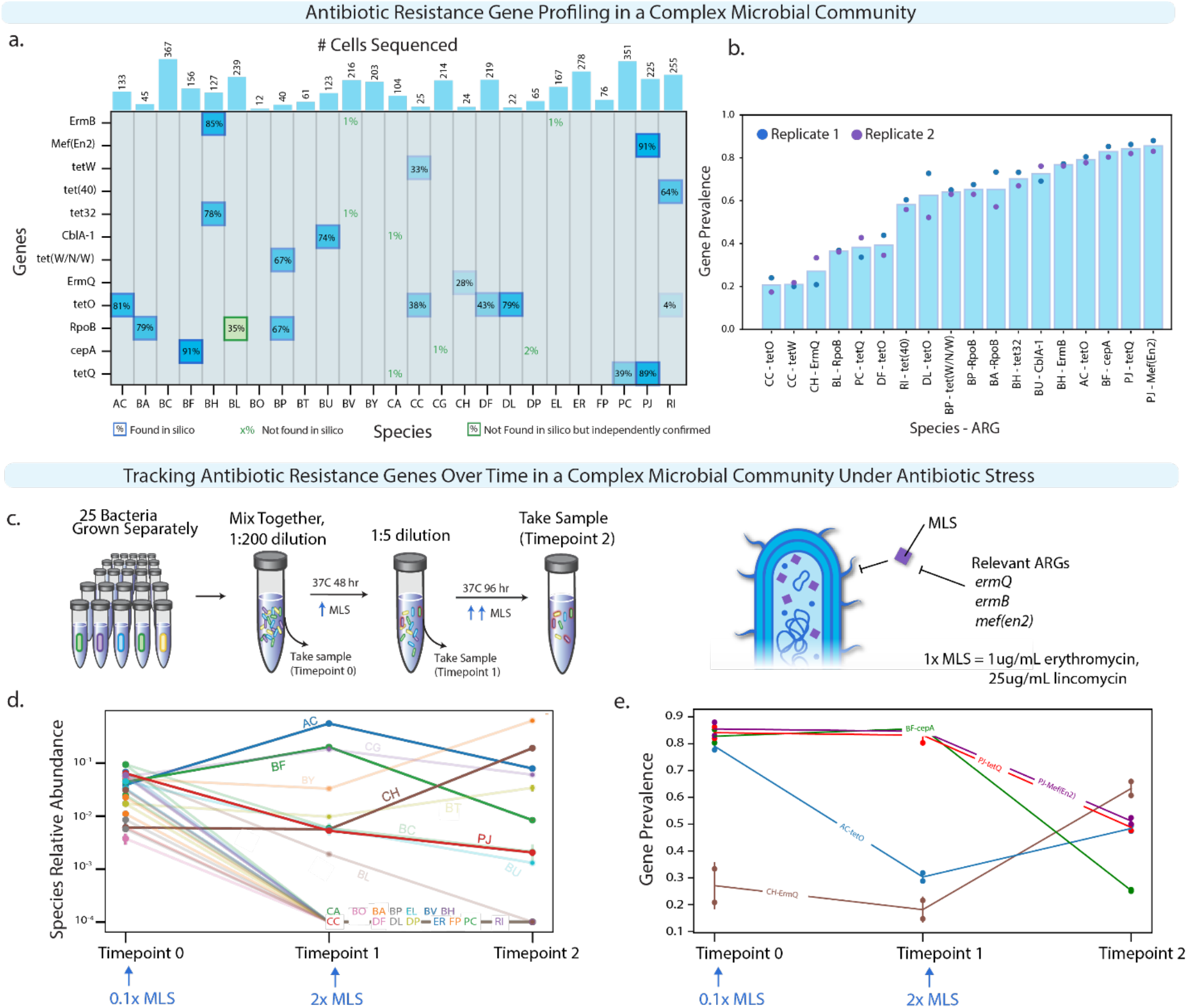
DoTA-seq enables tracking of gene-species dynamics in a complex human gut microbial community. **(a)** Heatmap of antibiotic resistance genes (ARGs)-species associations in a 25-member synthetic gut microbial community for a randomly chosen technical replicate. For each species (x-axis) and ARG (y-axis), the proportion of cells detected with the gene (prevalence) is shown. Bars on top show the number of cells sequenced for each species. The opacity of the background for each box is proportional to the prevalence value. Computationally predicted ARGs based on genome sequence are outlined in blue. ARGs that were not found in a given species’ genome sequence but observed using DoTA-seq as well as independently confirmed are outlined in green boxes. ARGs that were not found in a given species’ genome sequence are represented by green text with outline box. Species-ARGs combinations with a prevalence of less than 2% and where the gene is not found in the species’ genome sequence are likely due to background noise. **(b)** Bar plot of the average fraction of cells containing the ARG (i.e. gene prevalence) for each ARG-species combination that displayed greater than or equal to 10% prevalence. Data points represent technical replicates (n = 2). **(c)** Schematic of the experimental design for characterizing the ARG-species associations for a 25-member synthetic human gut community in response to antibiotics. The microbial community was cultured in the presence of increasing concentration of antibiotics. In this experiment, the community exposed to a lower concentration of antibiotic for 48 hours was passaged (i.e. community aliquot was transferred to fresh media) into media containing a higher concentration of antibiotic and then cultured for 96 hr. **(d)** Relative abundance of species at each measurement time determined by DoTA-seq. The lines corresponding to species that were not detected after passage 1 or did not contain ARGs are made transparent for de-emphasis. Error bars represent 1 s.d. from the mean of technical replicates (n = 2). **(e)** Prevalence of ARG-species associations at different measurement times for the antibiotic experiment. Species that were not detected after passage 1 and/or did not contain targeted ARGs are excluded from this graph. Data points represent technical replicates and error bars represent 1 s.d. from the mean of technical replicates (n = 2).

The measured prevalence is determined by both the proportion of cells containing the gene as well as by the differences in primer amplification efficiencies targeting the gene and the 16S primer (i.e. a low efficiency primer may skew measurements towards lower prevalence). To reduce these effects, relative amplification efficiencies between the different primer sets should be determined and adjusted based on analysis of the sequencing results (see **Supplementary Note 5**). Alternatively, gene prevalence could be compared between different conditions (i.e. relative changes) for a given gene-species pair, since potential primer biases are similar across the different conditions.

To investigate dynamic changes in gene prevalence, we cultured the 25-member community in the presence of erythromycin and lincomycin (MLS) at successively higher concentrations (**Fig. 3c**). The addition of MLS was expected to confer a fitness advantage to cells that harbored the *ermB, ermQ*, or *mef(en2)* gene. The DoTA-seq results showed that the growth of most species was inhibited in the presence of the antibiotics (**Fig. 3d**). Of the species that grew, the prevalence of several ARGs varied across timepoints (**Fig. 3e**). For example, in *Clostridium hiranonis* (CH), the prevalence of *ermQ* increased over time. In *Parabacteroides johnsonii* (PJ), the frequencies of *mef(en2)* and *tetQ* decreased. In AC, the prevalence of *tetO* decreased at the first timepoint, then moderately increased following the second timepoint.

To independently validate these trends, we performed colony PCR and single-cell PCR to independently assess the prevalence of *mef(en2), tetQ*, and *cepA* genes in gram-negative species PJ and BF from the samples described above (these techniques were difficult to perform on gram-positive cells). Consistent with DoTA-seq, single-cell PCR showed that *mef(en2)* and *tetQ* in PJ had frequencies of ∼0.8 at timepoint 0 and ∼0.6 at timepoint 2, while *cepA* had frequencies in BF of ∼0.7 at timepoint 0 and ∼0.2 at timepoint 2 (**Supplementary Fig. 7a**,**b**,**c**). Using selective plating of BF colonies, we further confirmed the decreasing trend of *cepA* prevalence in BF by PCR genotyping of colonies derived from passage 1 (7/14 colonies positive for *cepA*) and passage 2 (0/6 colonies positive for *cepA*) (**Supplementary Fig. 7d**). In sum, DoTA-seq revealed that ARGs can be present in variable fractions of bacterial populations in our conditions. In addition, the fraction of the population that harbors a given gene can drastically increase or decrease in response to antibiotic exposure.

### DoTA-seq reveals plasmid-host taxonomic relationships in a human fecal derived microbial community

To determine whether DoTA-seq could be applied to microbial communities derived from high complexity natural microbiome samples of unknown composition, we used DoTA-seq to characterize a natural microbial community derived from the ZymoBIOMICs fecal community standard **(Fig. 4a)**.

**Figure 4.**
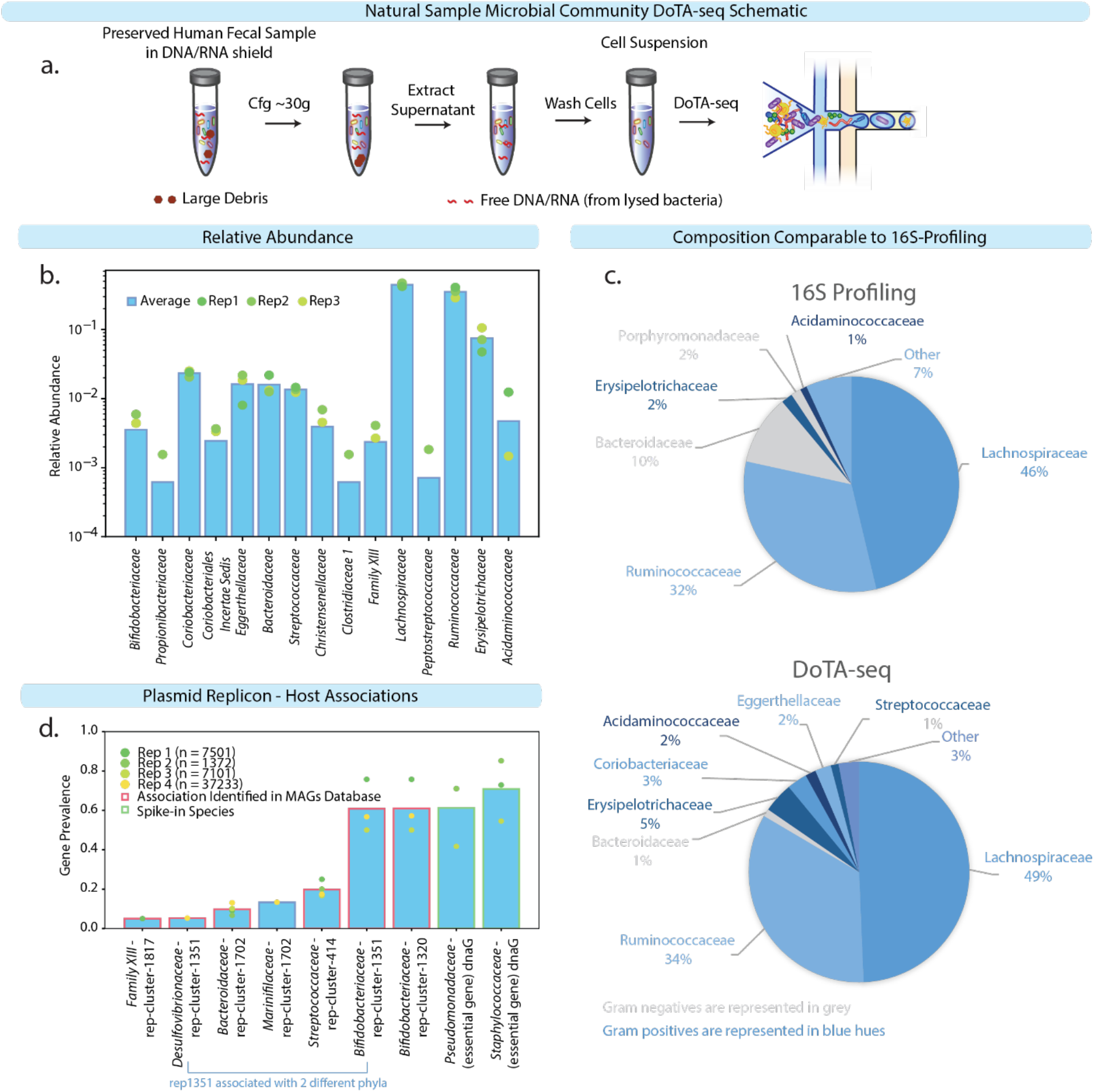
DoTA-seq can elucidate plasmid replicon-taxa associations in a fecal derived microbial community (ZymoBIOMICS human fecal standard). a) Schematic of experimental procedures. A human fecal sample preserved in DNA/RNA shield was gently centrifuged to remove large debris. The supernatant is extracted and washed to remove free DNA, leaving a suspension of cells, which was sequenced with DoTA-seq. b) Bar plot of the relative abundance at the family level. Each bar represents the mean and the markers represent individual replicates (n=3 technical replicates). The spike-in species are not included in the graph because they were at different relative abundances in each replicate (**Supplementary Figure 8**). C) Comparison of relative abundance obtained by 16S profiling (V3-V4 region) to DoTA-seq (V6-V8 region) of ∼37,000 cells. Gram-negative bacteria (Bacteroides for example) are under-represented in the DoTA-seq results, which is an artifact of the sample preservation method (DNA/RNA shield lyses gram-negatives) and thus not attributed to DoTA-seq **(Supplementary Fig. 2)**. d) Bar plot of the gene prevalence (the fraction of single cells of a given taxonomic identity containing a given plasmid replicon gene) of replicon-host associations (i.e. plasmid replication genes) detected in the DoTA-seq data. The bar represents the average of 3 replicates plus one additional replicate sequencing ∼37,000 cells. The circular markers represent individual replicates. The red outlines represent replicon-host associations that are also found in a database of human gut metagenome assembled genomes. The green outlines denote spike-in species that were not observed in the fecal derived community.

This human-associated fecal sample was preserved in DNA/RNA shield (https://www.zymoresearch.com/products/dna-rna-shield), which lyses gram-negative bacteria, and substantially reduces their ability to be captured by DoTA-seq **(Supplementary fig. 2)**. However, the gram-positive bacteria within this sample constitute a highly complex microbial community of unknown composition and thus can be used to evaluate DoTA-seq’s capabilities. To analyze single-cells within communities of unknown composition, we developed a novel data analysis pipeline (see Methods). We sequenced the community in three replicates of ∼3000 cells and a fourth library of ∼37,000 cells. The relative abundances of species were consistent across three replicates (**Fig 4b**). The taxonomic profiling from DoTA-seq with ∼37,000 cells displayed similar trends to 16S rRNA gene profiling **(Fig 4c**).

To assess the quantitative accuracy for both gram-negative and gram-positive bacteria, we introduced a spike-in of *P. putida* and *S. epidermidis* (species from two families not observed in the fecal sample). We spiked-in the two species at approximately equal number of cells (by hemacytometer counting) at three different abundances relative to the fecal community. DoTA-seq identified the relative ratio of *Pseudomonas* and *Staphylococcus* as 1.35 +/- 0.43 (mean +/- standard error) across the three different spike-in conditions **(Supplementary Table 2)**. This is reasonably close to the expected equal proportion since cell counting measurement errors are expected when generating the spike-in cells. The total number of observed spike-in cells decreased as their initial total abundance was reduced. In addition, DoTA-seq was able to reasonably recapitulate the quantitative changes in total relative abundance of the spike-in species to the fecal derived cells across conditions (**Supplementary Fig. 8**).

As a control for capture efficiency and cross-contamination rates, we targeted one essential gene (*dnaG*) in each spike-in species, which we expected in all spiked in cells and no other fecal derived cells. To identify plasmid replicon host associations, we targeted 9 plasmid replicon sequences identified in the metagenomic shotgun sequences of the fecal sample. Cross contamination of *dnaG* with the fecal community was not observed **(Supplementary Table 2**). In addition, we detected multiple associations between plasmid replicons and microbial taxa, including a replicon (rep 1351) that was associated with hosts from two different phylum in our sample (*Proteobacteria* and *Actinobacteria*) **(Fig. 4d, Supplementary Table 2)**. As an independent confirmation of these replicon-taxa associations, we performed a BLAST search of each plasmid replicon sequence to the metagenome assembled genomes (MAG) from the human microbiome for replicon-taxa associations with gene prevalence >= 5%. All observed replicon-taxa associations were present in the MAG database with the exception of *Marinfilaceae*-rep-cluster-1702 **(Fig. 4d, Supplementary Table 3)**. In sum, these results demonstrate that DoTA-seq is a powerful tool for studying the dynamics of mobile genetic elements and the dissemination of antibiotic resistance genes in complex natural microbial communities.

## DISCUSSION

Single-cell genetic heterogeneity underlies numerous important phenomena in the microbial world affecting evolutionary dynamics and microbiome functions^24,25,35,36^. Profiling genetic variation in complex microbial communities at the single-cell level is crucial for understanding them. However, major roadblocks to profiling single-cell genetic variation in microbiomes include efficient lysis conditions for diverse microbial species and efficiency of molecular biology reactions on low DNA template concentrations. To address these challenges, we developed DoTA-seq, a robust, accessible, and widely applicable tool for ultrahigh-throughput single-cell genetic profiling of microbes. Key to this method is cell encapsulation into reversibly crosslinked hydrogels with an appropriate pore size distribution. Encapsulation into hydrogels enables multi-step procedure for efficient lysis of diverse bacteria and simplification of microfluidics steps. We demonstrated the capability to profile genetic loci in up to ∼37,000 single-cells using 30 minutes of droplet making and one Miseq run. Therefore, it is reasonable to scale up to ∼3 hours of droplet making which corresponds to ∼10^5^ cells per run with an appropriate depth of sequencing. There are several aspects in DoTA-seq that can be tweaked to modify or improve performance. We discuss the most important ones below.

The encapsulation of unique barcoding primers at a limiting dilution captures ∼10% of the encapsulated cells. For many microbiomes and microbial populations, the number of available cells is not a limiting factor, and thus this capture frequency is acceptable. In cases where a large fraction of encapsulated cells need to be captured with unique barcodes, barcoded hydrogels^37^ could be used to replace the barcoded oligos. Hydrogels pack closely within the microfluidic device to enable delivery of a unique barcode to every droplet (instead of the ∼1 in 10 for limiting dilution), resulting in ∼100% cell capture. Alternatively, unique barcodes could be added at a higher encapsulation rate, such that on average, multiple unique barcodes are included per droplet, and very few droplets contain no barcodes at all. In this case, many cells would be represented by multiple unique barcodes according to the Poisson distribution. Downstream data analyses would have to take this additional layer into account by fitting a Poisson model to the barcode count data to infer the underlying single cell frequencies.

The use of microfluidics chips for droplet making presents a barrier to large-scale sample parallelization as compared to methods that can be performed in 96-well plates. However, that has not precluded the usefulness of chip-based library preparation techniques as exemplified by the success of commercial instruments like the 10X genomics Chromium^38^. Given that DoTA-seq requires only droplet making steps, a chip-less version of this workflow is theoretically possible using particle templated emulsions^39^, which could be developed in the future for those who require massive sample parallelization.

Designing specific primers that efficiently amplify the sequences of interest while minimizing off-target annealing is a crucial step in DoTA-seq. Primers that do not have these features can yield low rates of gene capture in DoTA-seq. The degree to which sequences are captured also depend on PCR conditions (e.g. type of DNA polymerase, buffer conditions, etc). Therefore, the absence of an amplicon can result from both the absence of a gene in the cell or a failure to capture a gene that is present due to low capture efficiency. We normalize the effect of capture efficiency by comparing gene frequencies relative to samples that are analyzed using the same protocol (i.e. primer sequences and reagents). For certain systems, we can overcome this limitation by designing primers that produce amplicons for all possible target gene states (e.g. unique amplicons for the ON and OFF state in the *B. fragilis* phase variation system), which enables absolute quantification.

The multiplex PCR primers used in DoTA-seq should be verified to not produce large primer dimers (“smears” larger than 300bp on a gel) when used together. Primer dimers that remain as <300bp fragments on a gel are typically tolerated as they can be removed using DNA size selection. In addition, each target should be amplified at similar efficiencies, though differences in the intrinsic amplification efficiency of primer sets can be compensated for by adjusting the relative primer concentrations. Although we have not explored the limits for the number of simultaneous targets possible in DoTA-seq, we generated a 12-plex DoTA-seq target assay with minimal effort using an automated script (see Methods: Generating DoTA-seq primers). In addition, commercial multiplex PCR amplification assays of up to ∼20,000 targets are reportedly available (CleanPlex, Paragon Genomics). This further supports the notion that a large number of loci could be simultaneously targeted with DoTA-seq. Previously characterized multiplex PCR primer sets, including well-characterized redundant PCR primer sets^40^ should perform well in DoTA-seq. Lastly, one pitfall to consider is that all cells regardless of their physiological or viability state are sequenced in DoTA-seq without discrimination. However, this pitfall is shared among all microbiome DNA-sequencing based methods and could be addressed using cell sorting to distinguish between different sub-populations based on fluorescent metabolic activity or viability dyes^41^.

The frequency of false positive (false association of a given gene with a taxonomic identity), serves as the limit of detection for rare events. These false positives arise from several potential factors including free DNA in the cell suspension, encapsulation of multiple cells per gel, diffusion of small DNA fragments (e.g. plasmids) between gels during storage, or infrequent merging of droplets during PCR. The likelihood of false positives is also determined by the abundance of the the target sequence within the sample. We have shown that the degree of false positives for small ∼4kb plasmids that are abundant in the population is ∼1%. Notably, this frequency of false positives is substantially lower for genetic loci harbored on bacterial chromosomes. To evaluate the false-detection rate, a target control gene (e.g. essential gene or a gene in a spike-in species containing a plasmid) could be analyzed across different experimental conditions.

DoTA-seq enabled us to profile the diverse subpopulations generated by phase variation of the CPS operons in the human gut symbiont *B. fragilis*. This new capability to observe combinatorial phase-variation enables future investigations of this system, which we explore in detail in another report^42^. Beyond the CPS operons of *B. fragilis*, mechanisms of rapid single-cell genomic variation, such as phase variation, antigenic variation^43^, and hypermutation^44^ are pervasive in microbes and play key roles in the lifecycles of pathogens. DoTA-seq will be a powerful tool for understanding these important phenomena.

DoTA-seq has a wide range of potential applications beyond the systems we explored here. For example, DoTA-seq could be used to quantify gene prevalence across different taxa, such as the genes for synthesis of the atherogenic compound trimethylamine in gut microbiota via a diverse set of metabolic pathways^40^, or the chemical conversion of cholesterol preventing its absorption in the gut^45^. Further, DoTA-seq can be used to interrogate the real-time microevolution of microbial populations by designing primers to target mutation hotspots^46,47^ determined from shotgun sequencing. The nucleotide sequence of these amplicons can be used to reconstruct phylogenetic lineages^19^. Furthermore, droplets can serve as highly parallelized compartments for culturing of small microbial communities^48^. In each droplet, DoTA-seq could be used to read out the composition of communities assembled in droplets by targeting the 16S rRNA gene of each species in ultrahigh-throughput. Finally, DoTA-seq could be adapted to probe the spatial heterogeneity of microbiomes via immobilization and fractionation in a gel matrix mirroring the MaPS-seq approach pioneered by Sheth et al^49^. In some applications, it is necessary to first characterize samples with metagenomics shotgun sequencing to identify genetic targets for subsequent single-cell analysis with DoTA-seq.

DoTA-seq represents a platform technology for building ultrahigh-throughput targeted single-cell analysis methods for diverse microbes. This generalizable and flexible workflow can be adapted to make other types of measurements in addition to sequencing of genetic loci. For example, performing DoTA-seq on cells pre-labelled with DNA barcode conjugated antibodies^50^ can enable single-cell profiling of targeted phenotypes such as protein expression, CPS expression, and other cellular phenotypes. In the future, we plan to expand this method to allow simultaneously profiling of genetic loci and target cell phenotypes via barcoded antibody labeling, generating genotype–phenotype matched single-cell data for microbial populations. This approach would further enrich the types of single-cell data available for microbiology and microbiome research.

## ACKNOWLEDGEMENTS

We thank Yu-Yu Cheng for providing the *E. coli* and *B. subtilis* strains and helpful discussions. We thank Ryan Clark for assistance with the synthetic human gut community. We are also grateful to Laurie Comstock for providing the *B. fragilis* recombinase deletion strain and Brian Pfleger for providing the *Pseudomonas putida KT2440* strain. This research was supported by the National Institutes of Allergy and Infectious Diseases under grant number R21AI156438, National Institute of General Medical Sciences under grant number R01GM38660 and R35GM124774.

## AUTHOR CONTRIBUTIONS

F.L. and O.S.V. conceived the study. F.L., O.S.V., J.S. and R.L. designed experiments and interpreted the data. F.L. performed experiments and analyzed data. J.S. designed the DoTA-seq assay and analyzed data for experiments involving phase variation in *B. fragilis*. T.D.R. designed scripts for analysis of single-cell digital PCR data. Z.Z. developed the data analysis pipeline for natural microbiome samples. F.L. and O.S.V. wrote the manuscript. F.L., O.S.V., R.L., K. A., and T.D.R. contributed to the revision of the manuscript.

## CONFLICTS OF INTEREST

F.L. and O.S.V. have filed a provisional patent Methods for isolating and barcoding nucleic acid. US Provisional Application 63/337468.

## DATA AVAILABILITY

Processed data containing barcodes and their mapped associated reads are available from Zenodo (https://doi.org/10.5281/zenodo.6537689). Raw sequencing data will be available from Zenodo at the time of publication.

## CODE AVAILABILITY

All code used in analysis of DoTA-seq sequencing data are available on Github: https://github.com/lanfreem/DoTA-seq-Paper.

## MATERIALS AND METHODS

### Oligonucleotides and primers

The oligonucleotides used were purchased from Integrated DNA Technologies as regular single stranded DNA oligonucleotides for universal sequences, an oligo-pool for the antibiotic resistance gene primers, and as Ultramers for the *B. fragilis* promoter primers. Ultramers were used to ensure higher synthesis yields. We found that best performance is achieved with individually ordered and validated primers and carefully combined manually to equal concentrations, rather than using the oligo-pool synthesis methods. Primer sequence and other details can be found in **Supplementary Tables**.

### Microfluidics fabrication

The master mold for the microfluidic devices was fabricated using soft lithography^51^ in a negative pressure cleanroom. Thin layers of SU-8 3025 photoresist (MicroChem) were applied to 3-inch silicon wafers (University Wafers) using a spin coating process to achieve layers of 20 μm for droplet generator 1 and 30 μm for droplet generator 2 respectively. Microfluidic feature patterns were then transferred to the SU-8 layers using a photolithography mask (CAD/Art Services) and a 365 nm collimated LED (Thorlabs M405L4-C1) at 120 mW for 1 minute and 45 seconds. Following exposure, the mold was soft-baked at 95 °C for 5 minutes before developing the patterned SU-8 in propylene glycol methyl ether acetate (PGMEA) for 2 minutes. The developed master mold was hard-baked at 200 °C for 2 minutes to complete curing of the SU-8.

Microfluidic devices were cast from the negative mold using polydimethylsiloxane (PDMS) (Sylgard-184) at a 1:11 crosslinker to elastomer ratio and cured at 65 °C for at least 12 hours. The devices were then cut from the master mold with a scalpel (Feather) and holes for the inlets and outlets were cut using a 0.75 mm biopsy punch (World Precision Instruments). To close the open microfluidic channels, a glass slide was bonded to the bottom of the PDMS devices. Chemical bonding between the PDMS and glass was achieved on contact following plasma activation with a plasma cleaner (Harrick Plasma). The completed microfluidic channels were treated with Aquapel Glass Treatment (Aquapel # 47100) to render the surfaces hydrophobic.

### DoTA-seq workflow

A cell suspension was stained with 1x SYBR Green (Invitrogen) and counted using a hemacytometer (Fisher scientific #0267151B) under a microscope to obtain the cell concentration. This concentration will be used to calculate the volume of cells to add to obtain the appropriate concentration for loading into the droplets (2.5 × 10^7^ cells/mL for lambda = 0.1). A polyacrylamide gel solution was prepared as follows: 100μL acrylamide monomer (Sigma-aldrich) in water (25% w/v), 15μL Bis-acryloyl Cystamine (Santacruz Biotech) in Methanol (Sigma-aldrich) (5% w/v), 10μL ammonium persulfate (Sigma-aldrich) (10% w/v), 75μL PBS, and the appropriate number of cells to obtain an encapsulation ratio of 0.1 in 4pL droplets. This solution was injected into a microfluidic drop maker (device 1) along with Biorad droplet generation oil for Evagreen (Biorad #1864005) with 0.5% v/v TEMED (Sigma) dissolved in the oil as catalyst. Droplet generation was carried out at 1000μL/hr for oil and 600μL/hr for the aqueous solutions. Collected droplets were incubated at 37°C for 10 mins for gel polymerization to complete. The polymerized gels were removed from the oil as follows: First the oil is drained using a pipette, and 1mL of acetone was added, then removed, then 1mL of isopropanol was added, then removed, then 1mL of PBS Wash Buffer was added to resuspend the gels in an aqueous buffer. The gels were then subject to 3 more washes in PBS Wash Buffer.

Cell lysis in the gels were carried out by adding 2x lysozyme buffer (20 mM Tris-HCl, pH 8.0; 10 mM EDTA; NaCl 100mM, 1% Triton X-100) with 20 mg/mL lysozyme (Sigma L6876) to equal volume of gels and incubating at 37°C for 30mins. The gels were then washed three times in 1mL PBS + 10mM EDTA, then added to equal volume of 2x proteinase K lysis buffer (Tris pH 8.0, 20mM EDTA, 100mM NaCl, 1% SDS) with 200 ug/mL proteinase K and incubating at 55°C for 30mins. The gels were finally washed four times in 1mL gel storage buffer (10mM HEPES pH 7.5, Tween 20 2%, NaCl 100mM, EDTA 20mM) and kept at 4°C until ready for use in barcoding.

Barcoding of the gels was carried out by first washing the gels four times in 1 mL pre-injection buffer (10mM HEPES pH 7.5, NaCl 25mM, EDTA 0.1mM, 2% Tween 20) achieving at least 1000x dilution. A PCR mix is made consisting of 1x Biorad ddPCR probes mix without dUTPs (Biorad 1863024), 400nM of P7, 40nM of Barrev-V3, 400nM of P5_I5_x where x represents the unique I5 index used for multiplexing libraries, 20nM (*E. coli/B. subtilis* and *B. fragilis* primers) or 5nM (ARGs primers) of each oligo in the targeted primer set, 0.015pM Barcode oligo (freshly diluted from 500pM stock), 2.5mM DTT (from single use aliquots) to a total of volume of 25μL. The gel and PCR mix are injected into microfluidic droplet maker device 2 (supplemental materials) with Biorad Evagreen droplet oil for encapsulation of gels with PCR mix using 200μL/hr for the gel and PCR mix, and 900μL/hr for the oil. Droplets are collected into a PCR tube (Fisher Scientific #14222292) until the PCR mix runs out. In the collected emulsion, excess oil is drained using a pipette, and thermocycled as follows: 95°C 5 min, 20 cycles of 95°C 30s, 72°C 10s, 60°C 5 min, 72°C 30s, then 20 cycles of 95°C 30s, 72°C 10s, 60°C 90s, 72°C 30s, then 72°C 10min. All steps except the first and last used 1°C/s ramp rate. For detailed video instructions on droplet making with gels, please refer to this video article by Demaree et al.^52^.

After PCR, the coalesced droplets were removed using a pipette, and the emulsion was broken on ice by adding 20μL 500mM EDTA and 20μL perfluoro-octanol (Sigma 370533), then vortexed followed by a spin pulse centrifugation. The aqueous phase was transferred to another tube by pipette, then 20μL 1M TCEP (UBP Bio P1021-100) was added to completely de-crosslink the gels, and the resulting solution vortexed for 10s to completely dissolve the gels. The whole mixture was cleaned up using a Zymo cleanup and concentrator kit (Zymo Research D4013), then subject to size selection using SPRI-select beads (Beckman Coulter B23317) using 0.7x volume of beads. A further round of size selection was performed in 100mM NaOH, 10% Ethanol, and 1.4x volume of beads to increase the purity of the library. For the most up to date and detailed DoTA-seq protocol, please refer to dx.doi.org/10.17504/protocols.io.n92ldzox7v5b/v1.

### Sequencing DoTA-seq libraries

The resultant libraries were quantified using qPCR (NEB E7630S) and sequenced on a Miseq (Illumina) using V3 150 cycles kit using custom read 1 and I7 primers. Up to 5 libraries were pooled per run.

### DoTA-seq of mixture culture of E. coli and B. subtilis

*E. coli* and *B. subtilis* strains (**Supplementary Table 4**) were grown overnight separately in LB broth (DOT scientific) with 34μg/mL of chloramphenicol and 5μg/mL of chloramphenicol, 1μg/mL of erythromycin, and 25μg/mL of lincomycin for *E. coli* and *B. subtilis*, respectively. 100μL of the respective overnight cultures were taken, mixed, then washed in 1mL PBS (Crystalgen) + 0.1% Tween-20 (Sigma-aldrich) twice before resuspending in 100μL of PBS. The cell suspension was then used as input for DoTA-seq using the *E. coli* and *B. subtilis* control primer sets (**Supplementary Table 5**).

### DoTA-seq of ZymoBIOMICS microbial community standard

For the microbial community standard experiments, 20μL of the ZymoBIOMICS Microbial Community Standard (Zymo #D6300) was washed twice in PBS + 1% Pluronic F68 (Thermofisher#24040032) and resuspended in 200 μL of PBS + 1% Pluronic F68. Cells were stained by incubation with 10X Cellbrite Fix 555 (Biotium #30088) and 10X SYBRGreen for 5 minutes and counted with a hemacytometer using CF555 fluorescence to estimate cell concentrations. These cells were prepared for sequencing using the Zymo Standard DoTA-seq primerset **(Supplementary table 6)** following the DoTA-seq V3 protocol which can be found at dx.doi.org/10.17504/protocols.io.n92ldzox7v5b/v1.

### Fluorescence microscopy of droplets and gels and counting of particles and droplets

Droplets (in oil) or gels (in aqueous buffer) were stained with SYBRGreen by mixing with equal volumes of SYBRGreen saturated Biorad Oil or dissolving SYBRGreen to 10x concentration, respectively. Next, 10μL of the SYBRGreen stained droplets or gels were pipetted into a disposable cell counting chamber slide (#C10228) and imaged on a Nikon Eclipse Ti epifluorescence microscope using 4x objective with a X-cite120 LED light source with 470/40nm filter and 560/40nm filters for SYBRGreen and CF-555 channels, respectively. The resulting raw images were analyzed using ImageJ (FIJI v1.53f51) using the Find Maxima function on the fluorescence and brightfield images to automatically count the numbers of fluorescent particles and droplets, respectively. In empty droplets or gels, the Find Maxima algorithm failed to identify fluorescent particles, as expected.

### DoTA-seq of the 25-member synthetic community

The 25 member synthetic community (**Supplementary table 1**) was prepared as in Clark et al.^33^. Briefly, 25 isolates were grown individually in an anaerobic chamber, then mixed to approximate equal proportions based on absorbance at 600 nm (OD600) using a Tecan F200 Plate Reader, with 200 μL in a 96-Well Microplate. Note that not all monocultures grew to sufficient OD to allow equal representation in the final community. The mixture of species was combined with 50% glycerol (Research Products International) and 400μL aliquots stored at -80°C. For each DoTA-seq experiment, a new glycerol stock was thawed, 200μL of cells were spun down and washed in 1mL PBS + 0.1% Tween-20 twice, before resuspension in 100μL PBS + 0.1% Tween-20. The cell suspension was used as input for DoTA-seq using the 25-member community primer sets **(Supplementary Table 7)**.

### DoTA-seq of the ZymoBIOMICS human fecal reference

The ZymoBIOMICS human fecal reference sample (Zymo D6323) was thawed at room temperature and 100 μL was taken and centrifuged at 35g for 20 minutes to separate cells from large fecal particles. The cell pellet containing supernatant was transferred to a new tube and washed twice with PBS + 1% Pluronic F68 and resuspended in 100 μL of PBS + 1% Pluronic F68. Next, 100 μL of overnight cultures of *Pseudomonas putida KT2440* and *Staphylococcus epidermidis ATCC 12228* grown in LB and BHI broth (Sigma # 53286) respectively were washed twice in PBS + 1% Pluronic F68 then resuspended in 100 μL of PBS + 1% Pluronic F68. All cells were stained with 10X SYBRGreen and counted on a hemacytometer to estimate cell concentrations. Fecal community and spike-in cells were combined at predetermined ratios and prepared for sequencing with the fecal sample DoTA-seq primer mix **(Supplementary Table 8)** using the DoTA-seq V3 protocol available on protocols.io at dx.doi.org/10.17504/protocols.io.n92ldzox7v5b/v1. For the large-scale experiment sequencing that captured >37,000 cells, the same protocol was followed except no spike-in cells were added and 250μL of the PCR master mix was used.

### Metagenomic shotgun sequencing and 16S profiling abundances for the ZymoBIOMICS microbial community standard and fecal standard communities

The raw reads and abundance data for both communities were taken from the respective datasheets of the products available at zymoresearch.com.

### Generating DoTA-seq target primers

For defined microbial communities, target genes were first identified from the genome sequences of the constituent strains. In particular, 100 candidate primers were generated from each gene using Primer3^53^ (http://primer3.org). The thermodynamic ΔG of primer self and heterodimerization of each candidate primer for all target primer genes were calculated using ntthal, which is a submodule of Primer3. A simulated annealing algorithm is then used to select primers for each gene that minimizes the potential of primer dimers. All steps are automated in Jupyter notebooks, which are available at Github (see Code Availability). For the fecal derived microbial community, the raw reads obtained from metagenomic shotgun sequencing (see above) are first assembled using Megahit v1.2.9^54^ using default options. The resulting contigs are searched using BLAST against the replicon database from Mob-suite^55^. We filtered the results to matches with more than 500bp in length and the top 10% identity. The plasmid replicon Inc18 was removed from the set of 10 targets because the primers generated large primer dimers in initial PCR tests. Since this primer set was not crucial for demonstration purposes, we simply removed this gene target from the list as opposed to testing different Inc18 primers.

### Sequencing the capsule polysaccharide (CPS) promoters in B. fragilis using DoTA-seq

*B. fragilis* DSM 2151 was streaked onto petri dishes containing Bacteroides minimal media with 1.5% w/v agar **(Supplementary Note 6)**. The plate was incubated in the anaerobic chamber at 37°C for 48 hours. After incubation, colonies were randomly picked using a pipette tip and resuspended into PBS + 0.1% Tween-20. The cell suspension was used as input for step 2 of the DoTA-seq using the *B. fragilis* CPS promoter primer sets, with a slight modification of the protocol. The first step of gel encapsulation and lysis was skipped, instead the cells were directly mixed with the PCR mix in the second step of DoTA-seq. The PCR mix used was NEB Q5 Ultra Mix (M0544S), and the thermocycling protocol was as follows: 98°C 2min, 40 cycles of 98°C 30s and 65°C 5 min, then 72°C 10min. Ramp rate was kept at 2°C/s. Primers used for these experiments are found in **Supplementary Table 9**.

### Culturing of the 25-member synthetic human gut microbial community

One glycerol stock of the 25-member synthetic community prepared as described above was resuspended in 2 mL of YBHI **(Supplementary Note 7**) in the anaerobic chamber and incubated at 37°C for 1 hour to recover. After one hour, the culture is split into two 1mL tubes for experimental replicates. YBHI is added to each tube to a total of 5mL with 0.1ug/mL of erythromycin and 0.25ug/mL of lincomycin. After 48 hours incubation at 37°C, 200μL of cells are taken out and added to 200μL of 50% glycerol, labeled “timepoint 1” and frozen down at -80°C. 1mL of the culture is added to 4mL of fresh YBHI with 2ug/mL erythromycin and 50ug/mL of lincomycin and incubated at 37°C anaerobically. After an additional 96 hours, 200μL of cells are taken out and added to 200μL of 50% glycerol, labeled “passage 2”, and frozen down at -80°C.

### Single-cell digital PCR

Cells derived from glycerol stocks of the synthetic human gut community exposed to antibiotics were thawed on ice, then 100μL of cells was washed with 1mL of PBS + 0.1% Tween-20, then resuspended in 30μL of PBS. 70μL of 100% Ethanol (Koptec) was added to fix the cells for 10 minutes at room temperature. After fixation, the cells are washed and then resuspended in 50μL of pre-injection buffer. A PCR mix was prepared using 10μL of Biorad digital pcr mix for probes, 1μL of 20X Primetime probe assay (IDT) for species specific *rpoB* gene, 1μL of 20X Primetime probe assay for the ARGs *cepA* for BF and *tetQ* and *mef(en2)* for PJ, 4μL of pre-injection buffer, and 4μL of washed cells in pre-injection buffer. 30μL of the Biorad droplet generation oil is added, and the solution is vortexed at the highest setting for 30s with a Biorad BR-2000 vortexer. The resulting emulsion is thermo-cycled for PCR as follows: 95°C 10min, and 40 cycles of 94°C 30s and 60°C 1min with a ramp rate of 2°C/s. 10μL of the resulting emulsion is loaded into a countess cell counting chamber (Invitrogen) and imaged with a Nikon Eclipse Ti epifluorescence microscope using 4x objective with a X-cite120 LED light source with 470/40nm filter and 560/40nm filters for FAM (*rpoB*) and HEX (ARGs) channels, respectively. For each sample, at least 4 images are counted in ImageJ to obtain the ratio of FAM/HEX positive droplets, using a custom macro script (see code availability) when more than 30 positive droplets are present and by manual inspection when less than 50 positive droplets are present. Primers for single-cell digital PCR are found in **Supplementary Table 10**.

### Colony PCR for B. fragilis and cepA

Cells derived from glycerol stocks from the synthetic human gut community antibiotic experiment were streaked onto Bacteroides minimal media agar plates and incubated for 48 hours in the anaerobic chamber for outgrowth of colonies. Individual colonies were picked using a pipette tip and inoculated into a YBHI growth media for growth overnight (∼16 hours) in the anaerobic chamber. Individual cultures were then spun down and resuspended in 1mL of TE buffer. A PCR mix consisting of 5μL ssoAdvanced Probes mix (Biorad), 0.5μL each of the 20X Primetime assay for *B. fragilis rpoB* and for *cepA*, 4μL of H20, and 1μL of the cell suspension. The mix was thermo-cycled in a Biorad CFX-connect real-time PCR system as follows: 95°C 3min, 40 cycles of 95°C 15s and 60°C 45s with fluorescence detection at 60°C. The samples with amplification detected in the HEX and FAM channels were determined to have originated from BF colonies and contain the *cepA* gene, respectively. *B. fragilis* was chosen for this experiment because a selective growth medium (Bacteroides minimal media, **Supplementary Note 6)** was available to isolate colonies from the community that were likely to be *B. fragilis*.

### DoTA-seq data analysis for synthetic communities

The raw sequencing reads are obtained from the Miseq. Reads 1 and 2 represent the targeted amplicons. The first index read represents the unique cell barcode while the second index read is used to multiplex libraries from different experiments. Demultiplexing of different libraries (index 2) is performed by the Miseq software. See **Supplementary Figure 9** for a flow chart of the analysis pipeline. Cell barcode demultiplexing and quality control is performed using a custom python script (R4-parser.ipynb). A typical library will yield ∼90% reads after filtering (see **Supplementary Table 11** for the sequencing performance of typical DoTA-seq libraries). The filtered reads were mapped to custom built reference databases containing *B. fragilis* CPS promoter sequences, 16S rRNA sequences and antibiotic resistance genes^56^ available on GitHub (see code availability) using Bowtie2 V2.3.4.1^57^ using “--very-sensitive” presets. A typical library will yield ∼90% mapped reads. The mapped reads were analyzed using custom python scripts as follows: The mapped reads are organized into read groups consisting of reads with the same unique cell barcode representing amplicons from the same droplet. Read groups with too few reads are removed. For each amplicon target for each read group, a minimum of 1% of the reads of the barcode group, or 5 reads, whichever is higher, is required to be present to count as a true “hit” for that target. This step removes background crosstalk between the barcodes in the library. When sequencing microbial communities containing different species, 16S rDNA amplicons within each read group are used to taxonomically identify the bacteria represented by the reads within the read group. Read groups with multiple 16S target matches are discarded. When sequencing *B. fragilis* CPS operons, only read groups containing amplicons for less than all 7 targeted amplicons or containing amplicons indicating conflicting promoter orientations are discarded. All scripts are available on GitHub (see code availability).

### DoTA-seq data analysis for natural microbial communities

Using DoTA-seq on communities of unknown composition presents unique data analysis opportunities and challenges. The ability to target multiple loci in a single cell presents the opportunity to simultaneously target multiple marker genes with short read sequencing to obtain more accurate taxonomic classifications. However, traditional pipelines for analysis of microbial taxonomic marker genes do not readily accommodate the single-cell barcoded data format. Marker genes originating from the same droplet should represent a single species. Leveraging this knowledge, we can first group marker gene reads by similarity for each droplet and build a consensus sequence for each group to correct for sequencing/PCR errors. We can then perform taxonomic classification for each droplet by using each consensus sequence for each droplet. We identify multi-encapsulated droplets as those that contain representative marker genes that represent distinct taxa. Thus far, we have implemented a proof-of-principle version of this analysis workflow for our natural microbial community datasets.

First, reads filtered for barcode quality score using standard DoTA-seq script (R4-parser.ipynb). Then, sequences were converted from “fastq” to “fasta” format with a minimum quality of Q20 using Seqtk (https://github.com/lh3/seqtk). Next, potential chimeras were filtered using USEARCH v11.0.667_i86linux32 using “silva_132_97_16S.fna” from the “SILVA_132_QIIME_release”^58^ as the reference using “sensitive” mode. Then OTUs for all chimera-filtered sequences were generated using MMseqs2 v13.45111^59^ with the settings of “-- min-seq-id 0.97 -c 0.95 --cov-mode 1”. Then, the OTU representative sequences were searched against “silva_132_99_16S.fna” from the “SILVA_132_QIIME_release”^58^ using BLASTN with the settings of “-evalue 0.001 -perc_identity 90 -max_target_seqs 1”. The final taxonomy hits were filtered by the criterion of sequence identity ≥ 95% and query coverage ≥ 90. Target genes (non-taxonomic markers) were identified using a database containing those genes and aligned using Bowtie2 as described in the standard DoTA-seq analysis workflow. The results of taxonomic classification of 16S OTU BLAST and target gene alignment with Bowtie2 are combined into one dataset and analyzed in a Jupyter notebook (Unknown-sample-analysis.ipynb). See **Supplementary Figure 9** for a flowchart depicting this pipeline.

### Identifying plasmid replicon sequences in MAGs

The website https://opendata.lifebit.ai/table/SGB contains 154,723 microbial genomes assembled from 9,428 samples of the human microbiome from diverse geographic locations, body sites, diseases, and lifestyles^60^. We downloaded all MAGs from the database, performed BLASTN search for all target replicon sequences with settings “-evalue 0.001 - perc_identity 0.9 -max_target_seqs 1, and filtered BLASTN results with an identity cutoff of 90%, replicon coverage cutoff of 70%. Within the filtered results, we used GTDB-Tk^61^ v1.8 to extract taxonomic information for all selected MAGs, which contained hits to the replicon sequences.

