## Supplementary Information for "Massively parallel single-cell sequencing of genetic loci in diverse microbial populations"

**Rep1**

| Species | GFP | dnaPolIII-Beta | RFP | dnaG | # cells |
|---|---|---|---|---|---|
| *B. subtilis* | 0.928 | 0.931 | 0.010 | 0.003 | 2639 |
| *E. coli* | 0.003 | 0.001 | 0.970 | 0.910 | 10752 |

(Columns GFP through dnaG: Frequency Found in Cells)

**Rep2**

| Species | GFP | dnaPolIII-Beta | RFP | dnaG | # cells |
|---|---|---|---|---|---|
| *B. subtilis* | 0.859 | 0.871 | 0.010 | 0.003 | 2311 |
| *E. coli* | 0.002 | 0.001 | 0.950 | 0.898 | 7930 |

(Columns GFP through dnaG: Frequency Found in Cells)

**Supplementary Figure 1.** Table of gene frequencies in *B. subtilis* and *E. coli* cells in the control experiment (Figure 1). Frequencies were computed by dividing the number of cells of a given species harboring a given gene divided by the total number of cells. GFP and dnaPolIII-Beta (in green) should be found only in *B. subtilis*, and RFP and dnaG (in red) should be found only in *E. coli*.

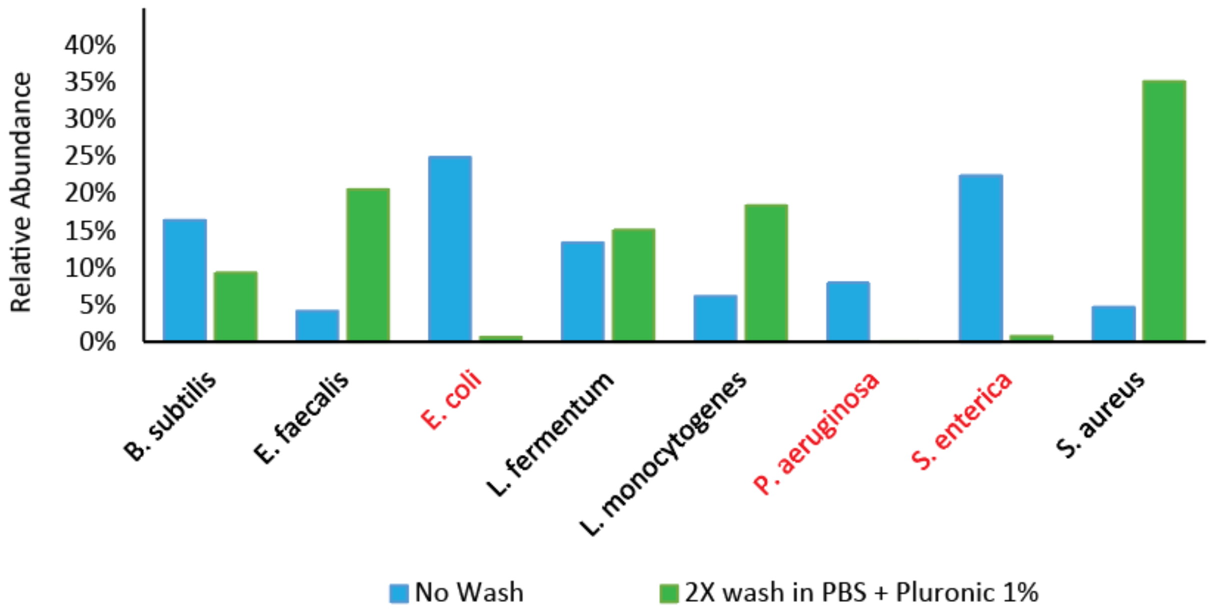


**Supplementary Figure 2.** DoTA-seq relative abundance of a Zymobiomics microbial community standard preserved in DNA/RNA shield with and without washing the cells. Gram-negative cells (in red) are almost undetected after 2X wash. This suggests that gram-negative cells were already lysed in the buffer and washing removed the cell debris and free-floating genomic DNA, leaving only the gram-positive cells for DoTA-seq.

**a**

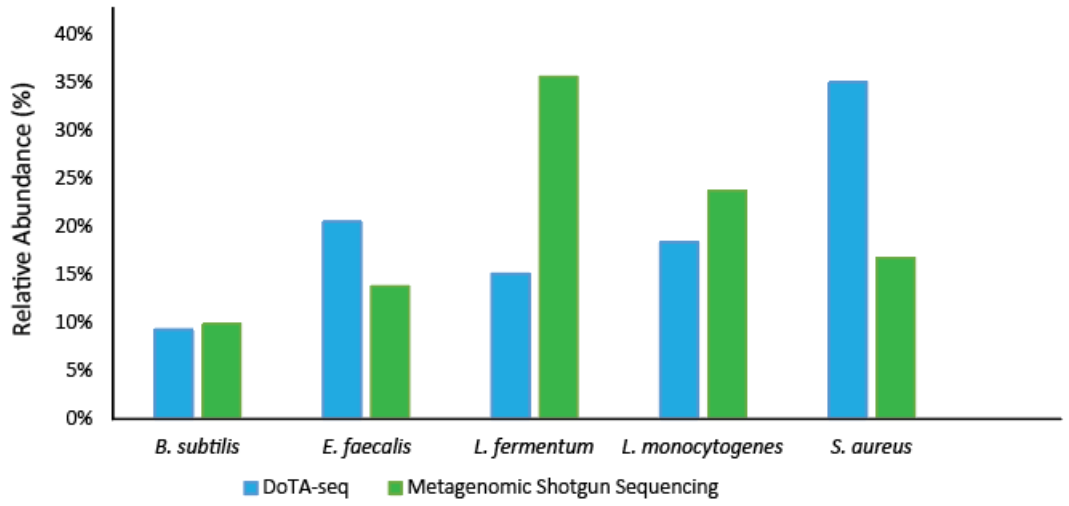


**b**

**Essential Gene *RpoB* (named by species of origin)**

| Species | *Bacillus subtilis* | *Enterococcus faecalis* | *Escherichia coli* | *Lactobacillus fermentum* | *Listeria monocytogenes* | *Pseudomonas aeruginosa* | *Salmonella enterica* | *Staphylococcus aureus* | Number of cells |
|---|---|---|---|---|---|---|---|---|---|
| *Bacillus subtilis* | 72% | 1% | 0% | 0% | 0% | 0% | 0% | 0% | 209 |
| *Enterococcus faecalis* | 0% | 71% | 0% | 0% | 0% | 0% | 0% | 0% | 482 |
| *Escherichia coli* | 0% | 0% | 0% | 0% | 0% | 0% | 0% | 0% | 16 |
| *Lactobacillus fermentum* | 0% | 0% | 0% | 92% | 0% | 0% | 0% | 0% | 351 |
| *Listeria monocytogenes* | 0% | 0% | 0% | 0% | 64% | 0% | 0% | 0% | 431 |
| *Pseudomonas aeruginosa* | 0% | 0% | 0% | 0% | 0% | 0% | 0% | 0% | 1 |
| *Salmonella enterica* | 0% | 0% | 0% | 0% | 0% | 0% | 10% | 0% | 21 |
| *Staphylococcus aureus* | 0% | 1% | 0% | 0% | 0% | 0% | 0% | 76% | 801 |

**Supplementary Figure 3**. DoTA-seq results of the Zymobiomics microbial community standard. **(a)** Comparison of relative abundances for the Gram-positive species obtained by DoTA-seq and by metagenomic shotgun sequencing. Results are mostly similar with the exception of some over-representation of Staphylococci and Enterococci in the DoTA-seq results. **(b)** Gene prevalence of essential genes captured by DoTA-seq in the microbial community. Intensity of cell color highlighting (green) is proportional to the prevalence for that gene-species combination. Gram-positive bacteria show high capture rates suggesting successful capture of whole-cells in gels, compared to Gram-negative bacteria which show low abundance and low capture rate of essential genes, suggesting that the initial sample did not contain whole un-lysed cells that could be captured by the gels. The gene prevalence are on average lower than that of the *Bacillus-E. coli* control experiment because the primer-ratios have not been optimized. We expect that optimization of the gene target primer ratios will improve capture efficiencies.

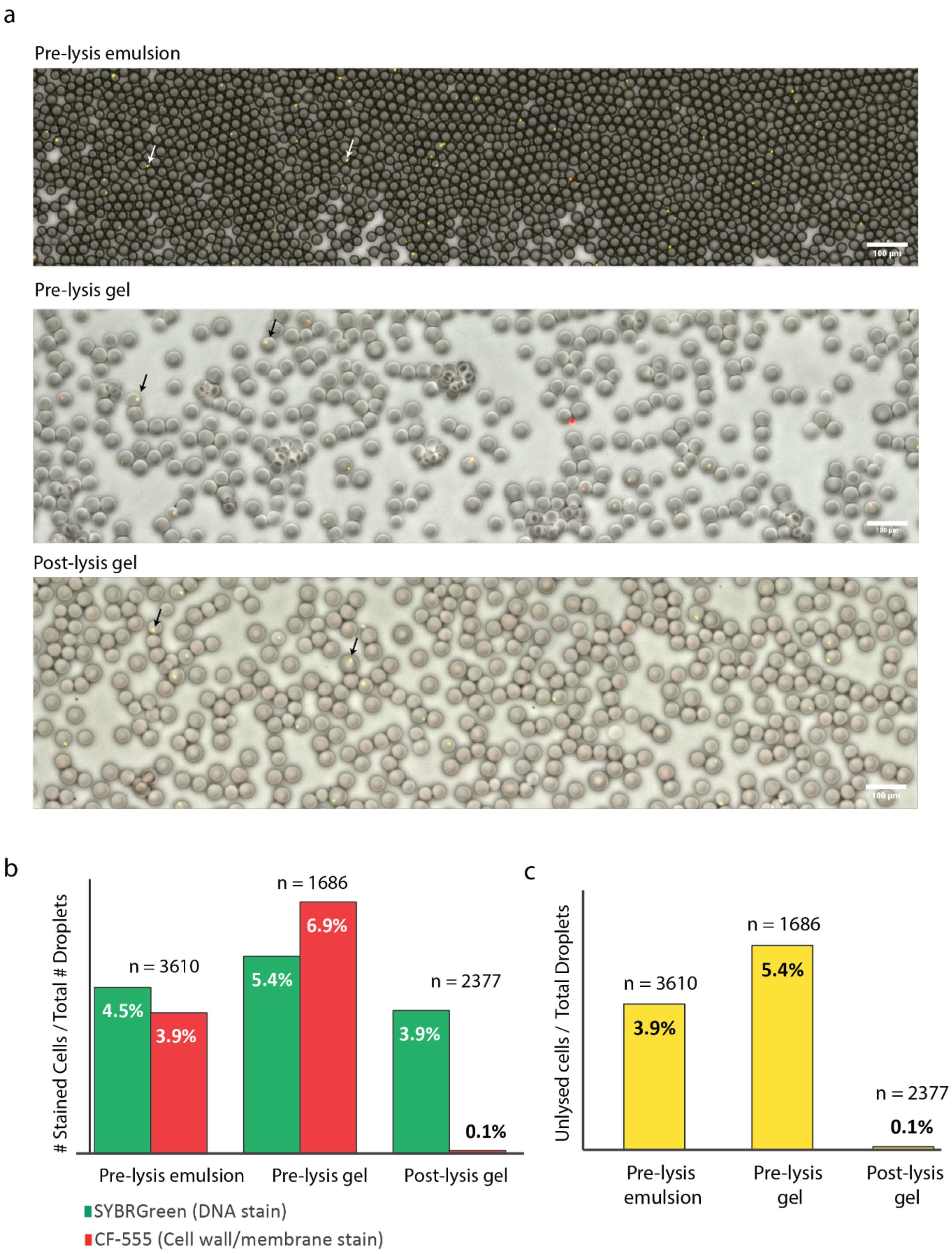
a
Pre-lysis emulsion
100 µm
Pre-lysis gel
Post-lysis gel
100 µm
b
### Stained Cells / Total # Droplets
n = 3610
n = 1686
n = 2377
4.5%
3.9%
5.4%
6.9%
3.9%
0.1%
Pre-lysis emulsion
Pre-lysis gel
Post-lysis gel
SYBRGreen (DNA stain)
CF-555 (Cell wall/membrane stain)
c
Unlysed cells / Total Droplets
n = 3610
n = 1686
n = 2377
3.9%
5.4%
0.1%
Pre-lysis emulsion
Pre-lysis gel
Post-lysis gel

**Supplementary Figure 4.** Tracking cell lysis efficiencies using Cellbrite-Fix 555 and SYBR-Green staining in >1000 droplets or gels in each step of the lysis protocol. Cellbrite (CF555) stains cell membrane and cell wall components while SYBR-Green stains DNA, allowing us to identify whether a cell is intact (CF555 and SYBR staining) or lysed (only SYBR staining). **(a)** Representative fluorescence microscopy overlay of Green Channel (SYBR staining), Red Channel (CF555 staining), and Brightfield (Droplets/Gels) as a water-in-oil emulsion, as gels before lysis (top two images) and as gels after lysis (bottom image). Arrows denote representative droplets containing cells. Scale bars represent 100μm. **(b)** The percentage of droplets containing fluorescent particles (# of fluorescent particles divided by the total # of droplets or gels) by SYBR or CF555 staining for each condition. Before lysis, CF555 and SYBR fluorescent particles are approximately equal in abundance. Following lysis, CF555 particles are substantially reduce, indicating lysis of the cells. The variable n represents total number of droplets analyzed. **(c)** The percentage of droplets containing unlysed cells at each step. Unlysed cells are defined as particles that are fluorescent in both CF555 and SYBR Green channels.

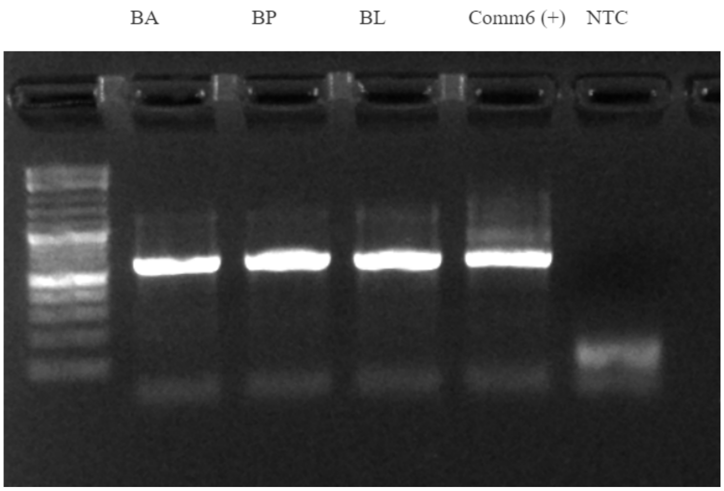


**Supplementary Figure 5.** Agarose gel showing PCR amplification of the *Bifidobacterium* specific *rpoB* gene on the genomic DNA extracted from pure cultures of *B. adolescentis* (BA), *B. psuedocatenulatum* (BP), *B. longum* (BL), or the 25-member synthetic human gut community. NTC represents no template control.

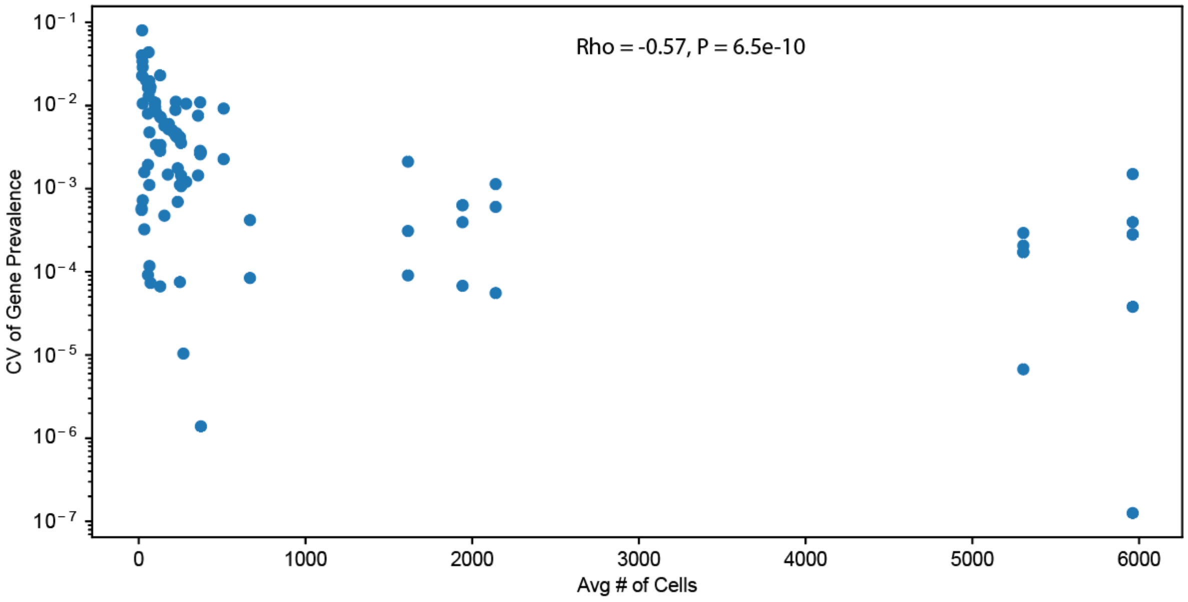


**Supplementary Figure 6.** Scatter plot of the average number of cells for that species versus the coefficient of variation (CV) of gene prevalence for two technical replicates for every gene-species pair. CV is calculated as the standard deviation divided by the mean. The negative Spearman correlation (Rho) suggests that a moderate fraction of the variance found between technical replicates can be attributed to stochasticity due to small numbers of cells.

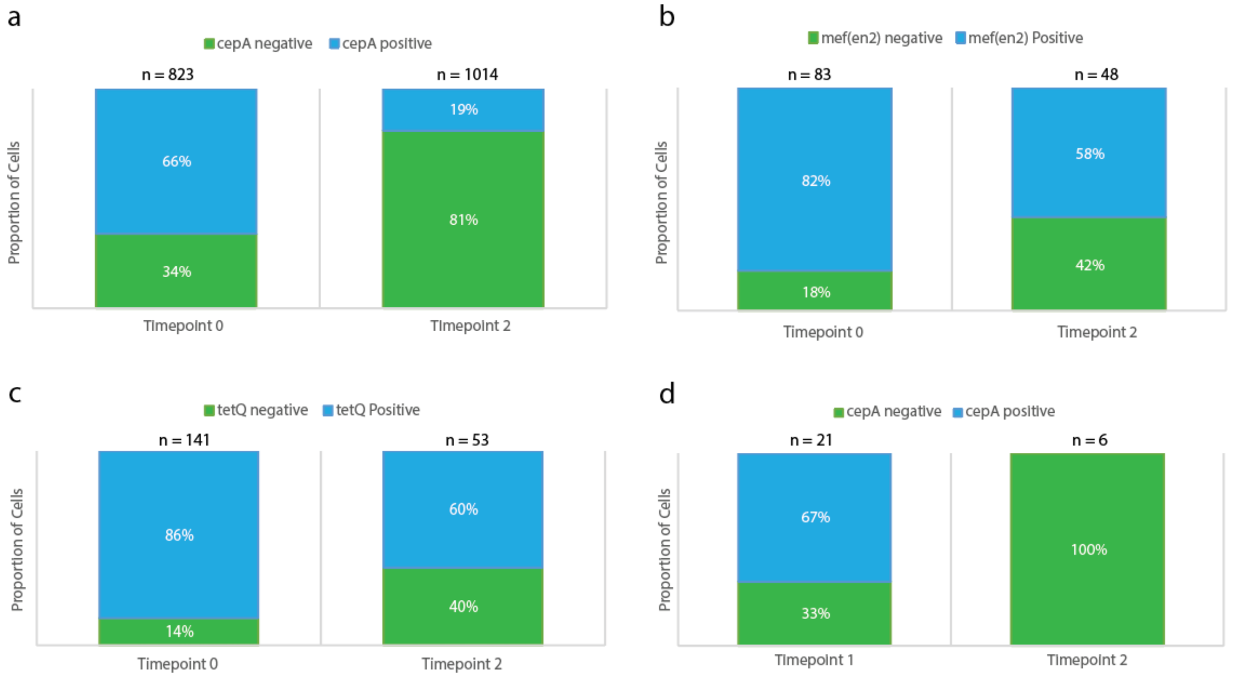


**Supplementary Figure 7.** PCR genotyping of cells derived from culturing of the 25-member synthetic gut community in the presence of antibiotics for samples at two timepoints. Droplet PCR genotyping for **(a)** *B. fragilis* (BF) and the *cepA* gene, **(b)** *P. johnsonii* (PJ) and the *mef(en2)* gene, or **(c)** PJ and the *tetQ* gene. Cells were fixed and subjected to digital PCR amplification for species marker genes and ARGs in an emulsion PCR (see Methods). Blue bars show the proportion of droplets that showed amplification for the species marker gene as well as the ARG, green bars show the proportion of droplets that show amplification for the species marker gene (*rpoB*) but not the ARG. Time 0 and time 2 represents timepoints 0 and 2 from the synthetic community antibiotic experiment, respectively. n represents the number of species marker gene positive droplets that were counted in total for each condition. **(d)** Colony PCR genotyping of *B. fragilis* (BF) colonies of cells originating from the synthetic human gut community experiment exposed to antibiotics. Colonies from glycerol stocks of samples taken at timepoints 1 and 2 of the synthetic community antibiotics experiment shown in Figure 3 are genotyped by qPCR for the BF specific *rpoB* gene and the antibiotic resistance gene *cepA*. The proportion of *cepA* positive colonies are shown in red, and proportion of *cepA* negative colonies are shown in blue. n represents the number of colonies that were positively identified as BF by successful amplification of the BF specific *rpoB* gene. In total, 24 colonies from timepoint 1 and 32 colonies from timepoint 2 were genotyped (many colonies were not BF).

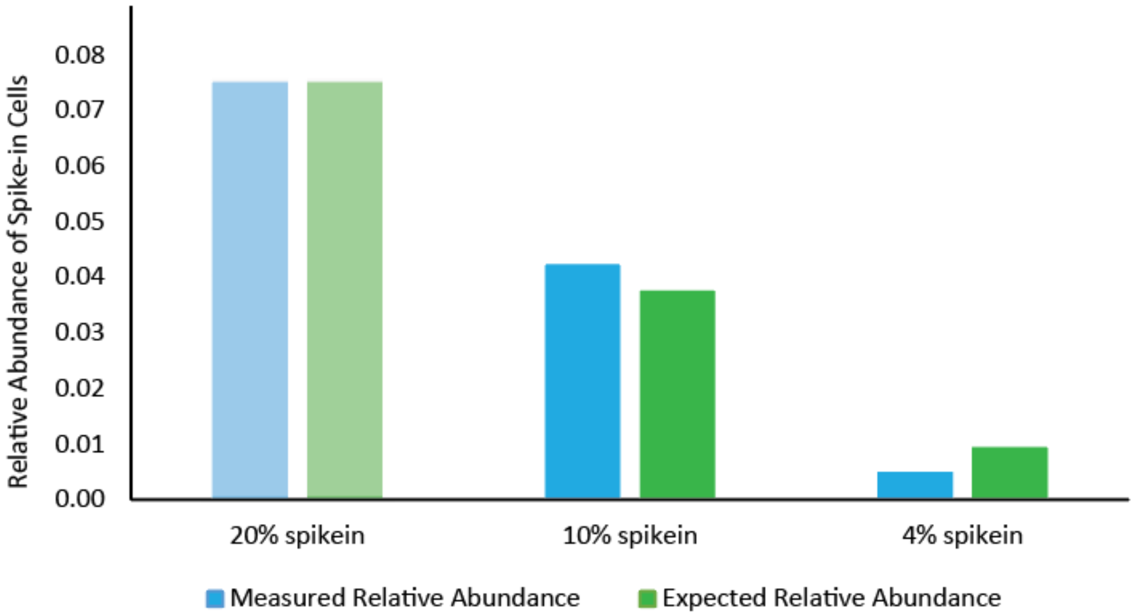


**Supplementary Figure 8.** The measured and expected relative abundance of spike-in cells (*Staphylococcus epidermidis* and *Pseudomonas putida* combined) for varying abundance of spike-in cells used in sequencing the fecal derived sample libraries. Since the theoretical number of cells in the fecal sample is not known a priori, the expected relative abundance is normalized to be the measured relative abundance for the 20% spike-in condition.

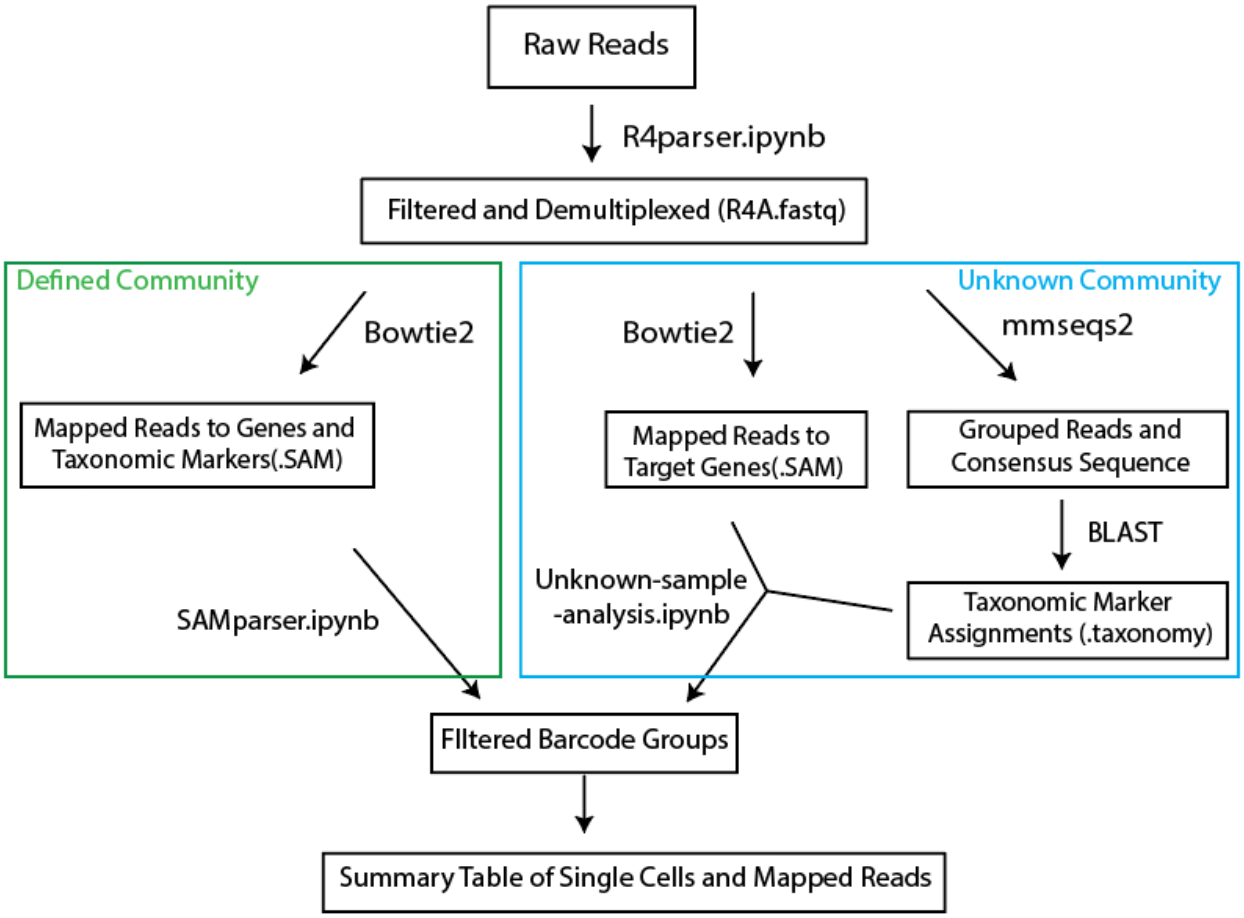


**Supplementary Figure 9.** Sequence analysis and filtering pipeline for DoTA-seq. Boxed text represent data and arrows represent scripts or programs used to process the data. Green box contains analysis specific to communities of known composition, and blue box contains analysis specific to communities of unknown composition. Raw reads obtained from the sequencer are processed in Python (R4parser.ipynb) to remove low quality barcodes and associate the droplet barcodes with the amplicon reads. The resulting reads are processed differently depending on the source sample material. For defined microbial communities, reads are mapped to a reference containing all target genes and taxonomic marker genes using Bowtie2, obtaining a SAM file. This SAM file is analyzed using Python (SAMparser.ipynb) to filter droplet barcodes based on evidence of cross-contamination and minimum numbers of reads (more details in the Github README file). For reads from unknown (i.e. naturally derived) communities, all reads are first mapped to a reference containing the non-taxonomic marker target genes using Bowtie2. For taxonomic classification, all reads are first clustered by pairwise similarity using mmseqs2, then the consensus sequence for each grouped cluster is used to search a BLAST database of taxonomic markers (such as 16S rRNA database). These results are combined with the Bowtie2 results then filtered based on markers of cross-contamination and minimum numbers of reads using Python (Unknown-sample-analysis.ipynb). More details are available in the README file on the Github page.

**Supplementary Note 1:** Poisson statistics for DoTA-seq

To achieve single-cell sequencing in DoTA-seq, limiting dilution is used to ensure that most droplets contain at most one cell and one unique DNA barcode. Assuming uniformity of droplet sizes, the distribution of cells or barcodes per droplet at limiting dilution follows the Poisson distribution:

$$P(k) = \lambda^k e^{-\lambda}/k!$$

Where $\lambda$ represents the average number of cells or barcodes expected per droplet, and $k$ represents the number of cells or barcodes per droplet. For $\lambda = 0.1$, the proportion of droplets expected to contain greater than one cell or barcode $P(k > 1)$ is 0.00468 or ~0.5% of the droplets. On the other hand, the fraction of empty droplets $P(k = 0)$ is 0.905 or ~90% of the droplets. The fraction of droplets containing exactly one cell or barcode is $P(k = 1)$ is 0.090 or ~ 9%.

As a result, assuming that we use a $\lambda = 0.1$ for both cell encapsulation in step 1 and barcode encapsulation in step 2, we will expect that the majority of the droplets will contain no cell or barcode. We would expect that $P(k = 1)$ for cells x $P(k = 1)$ for barcodes = ~9% x 9% = ~8% of total droplets will contain exactly one cell and one barcode.

**Supplementary Note 2:** Filtering out barcodes containing multiple cells through unique sequence signatures

Multiple cells can be tagged by a single barcode through multiple processes. Here they are in increasing order of relevance:

1. The probability of two randomly sampled barcodes containing the same sequence is exceedingly rare and is not expected to contribute noticeably to the data generated.
2. During the second droplet making step of DoTA-seq, two gels can sometimes be encapsulated into a single droplet when the frequency of droplet making does not perfectly match the frequency of gel reinjection. The rate of this happening depends on how well the microfluidics device is functioning, which is dependent on quality of fabrication and mono-dispersity of the gels. For a well-functioning device with high quality gels, this effect is negligible. The extent to which this is happening can be estimated by looking at the emulsion under the microscope post encapsulation. Droplets containing multiple gels will appear larger and irregularly sized under the microscope.
3. The probability of randomly encapsulating two cells in a single gel during the first encapsulation step is described by the Poisson distribution. For a Poisson loading ratio of 0.1 cells per droplet, the expected rate of two or more cells per droplets approximately 5% of all droplets that contain cells. We can confirm the expected rate of multi-encapsulated gels through fluorescence microscopy of SYBRGreen stained cells.
4. Droplets may coalesce during handling of emulsion and PCR thermocycling, which may result in droplets containing different cells merging into a single droplet, thereby producing barcodes that simultaneously tag multiple cells. One can determine the extent of coalescence by viewing the emulsion under the microscope after PCR. Improvements in surfactant chemistry will reduce the incidence of droplet coalescence during PCR. Droplet coalescence can also be detected by looking at the sequencing data **(Fig. SN1)**. Plotting the normalized cumulative fraction of reads against the number of reads per barcode reveals barcodes that contain more reads than expected. These likely resulted from coalesced PCR droplets which are bigger and produce more reads on average than a normal sized droplet.

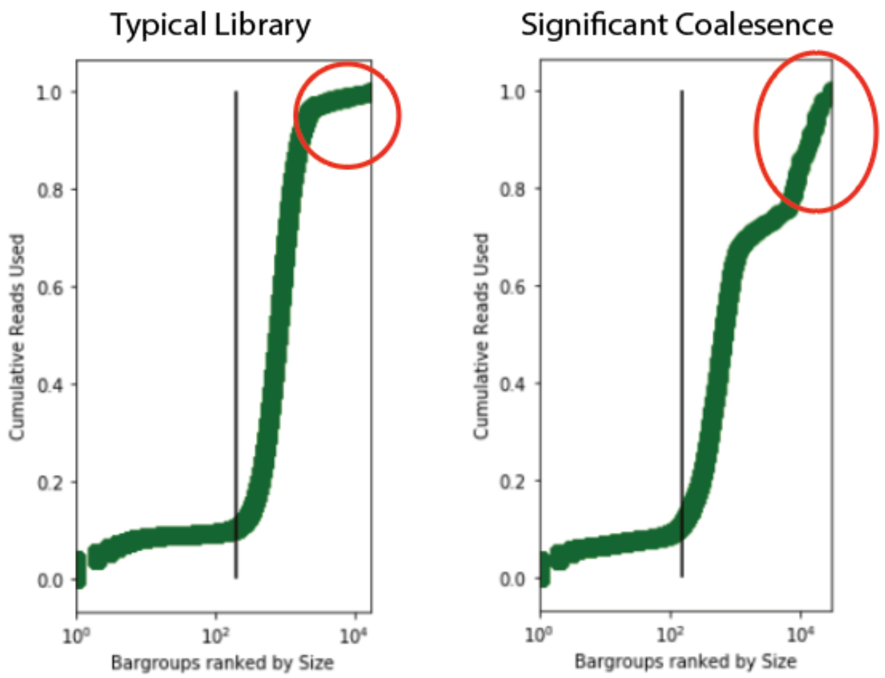


**Figure SN1** Cumulative reads plotted against the barcode group size (number of reads tagged by a unique barcode) revealing barcodes from coalesced droplets (circled in red). Left: the typical library has a very small population of coalesced droplets (accounts for very little cumulative reads). Right: a library with a large proportion of coalesced droplets (accounting for >20% of cumulative reads). Vertical line represents the cutoff where barcodes to the left represent mutated or erroneous barcodes that are removed from analysis (represents <10% of cumulative reads).

Usually, these quality control steps are not crucial as the resulting barcodes that tag multiple cells can be filtered out in data processing. To filter out barcodes that have tagged multiple cells, we look for signatures of conflicting reads in the sequencing data that indicates the presence of multiple cells. For multi-species populations, taxonomic marker genes such as the 16S rRNA will contain reads that represent different species, which cannot come from a single cell. These can be filtered out. For populations that contain a single species, if sequencing reads from the same locus contains conflicting sequences (for example, promoter in both the On and Off orientation) indicate that a single barcode has tagged multiple cells. Using these methods, we typically remove ~10% to~ 30% of our barcodes from our libraries.

However, this method does not account for cases where the same barcode tags multiple cells with the indistinguishable genetic sequences at the targeted (e.g. single population, no genetic sequence variation). If such cases are expected to significantly affect experimental results, we recommend strictly following the quality control measures described above (1-4) to minimize the likelihood of these events.

**Supplementary Note 3:** Representative commercial vendors for droplet making microfluidics equipment

Dolomite Microfluidics: https://www.dolomite-microfluidics.com/microfluidic-systems/single-emulsions/

Elve Flow: https://www.elveflow.com/microfluidics-application-packs/microfluidics-packs/easy-droplet-generation/

Chipshop: https://www.microfluidic-chipshop.com/catalogue/microfluidic-chips/polymer-chips/droplet-generator-chips/

Precigenome: https://www.precigenome.com/microfluidic-droplet-generator

**Supplementary Note 4:** Stochastic limit of detection for rare populations/variants and considerations for number of cells to sequence per sample.

The probability of detecting k cells from a population with relative frequency *p* by randomly sampling *n* cells is described by the binomial distribution:

$$P(k,p) = \binom{n}{k} p^k (1-p)^{(n-k)}$$

For the *B. fragilis* capsule subpopulations, the probability of detecting at least one cell is equal to the probability of not detecting any cells P(k > 0, *p*) = 1 – P(k = 0, *p*). We set the stochastic limit of detection to the frequency *p* where we would detect at least one cell 80% of the time P(k>0, *p*) = 0.8.

This equation simplifies to

$$P(k > 0, p) = 1 - (1-p)^n = 0.8$$

For n = 5000 cells, we calculate p to be approximately 3e-4.

More generally, the probability of detecting at least m cells from a population of frequency p in the sample is equal to P(k ≥ m, p) = 1 – P(k < m, p). These probabilities can be solved with a computer to derive the probability of seeing a rare population with frequency *p,* m times, for n cells sequenced in total.  An example of the results of this calculation is provided below for *p = (0.1, 0.01, and 0.001)* and m = 100 (To detect at least 100 cells of a specific type that is present at 10% – 0.1% relative abundance).

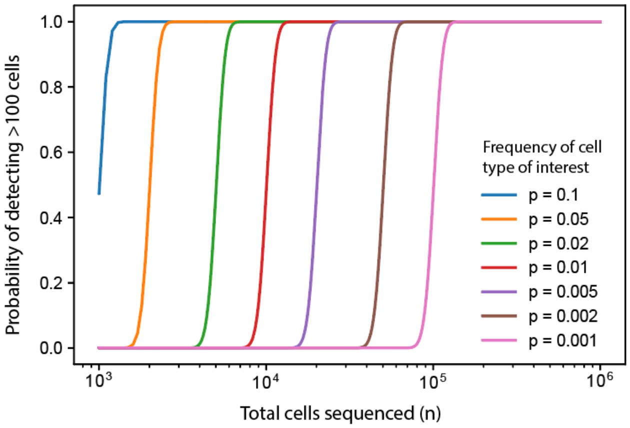

**Supplementary Note 5: Determining relative capture efficiencies for target primer versus 16S primer using sequencing data.**

Differences in primer amplification efficiencies can translate to different ratios of resulting amplicons within each droplet. In the extreme case, a droplet could contain mostly the 16S amplicon and the target gene amplicon could be missed by sequencing despite it being present on the cell genome. To avoid this phenomenon, we must ensure that the capture efficiencies for each target primer is reasonably balanced with other primers that we expect to co-amplify within the same droplet. In most cases, the other amplicon is the 16S amplicon.

To determine the relative capture efficiencies between the 16S amplicon and the target primer, we perform a DoTA-seq run using a sample that we know to contain cells with the target genes. Analyzing the sequencing reads, we tabulate the ratio of the target reads to 16S reads for each set of reads representing a single droplet. Plotting these ratios for each target primer, we will discover a distribution of target/16S ratio of the resulting reads. Below is an example of a set of distributions for 3 plasmids that were targeted using DoTA-seq in an initial pilot run.

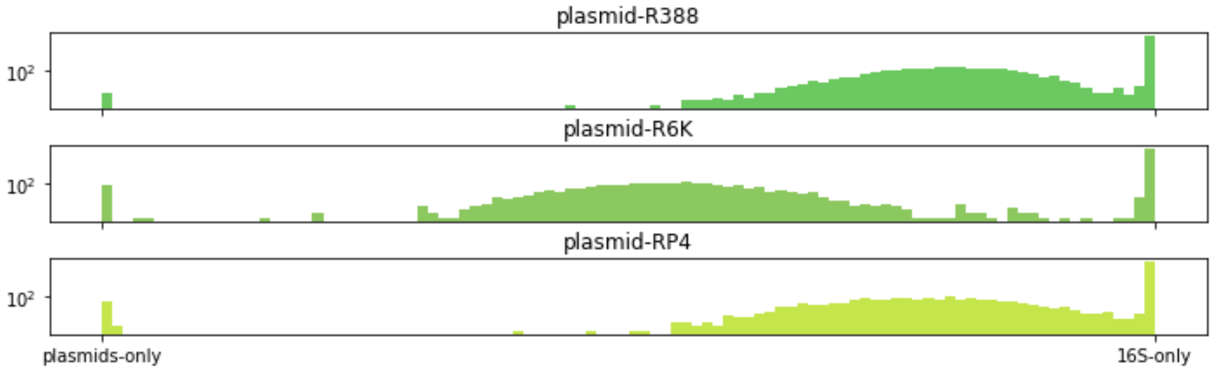


Plasmid R388 and plasmid RP4 distribution skews towards 16S amplicons, which means that the capture efficiency for the target primers are lower than that of 16S. Therefore, we can adjust the primer concentrations for the R388 and RP4 target primers accordingly to result in more balanced capture efficiencies (see below).

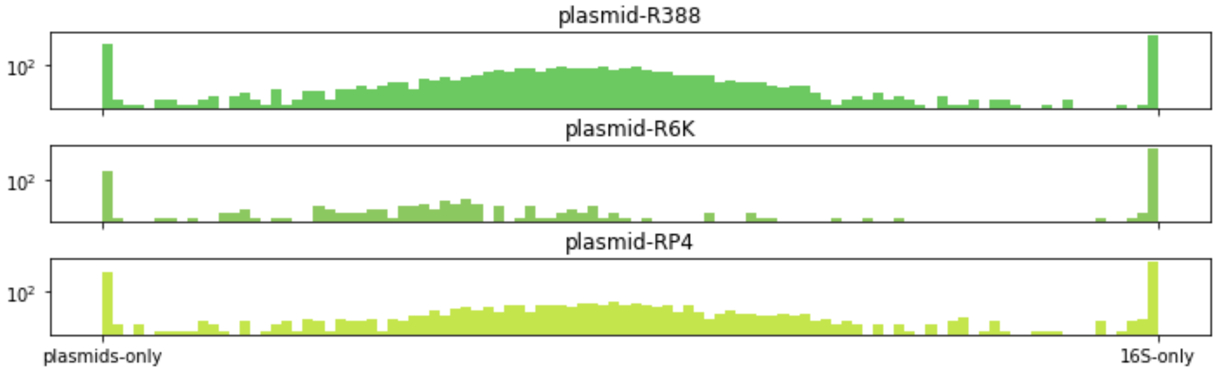

Sometimes it’s desirable to even skew the distribution towards the target genes so that we maximize gene capture at the cost of throwing aways more cells that do not have any 16S reads. It is impossible to always capture both the 16S amplicon and target amplicon in all cases, but adjusting the primer ratios will serve to maximize the effectiveness of DoTA-seq.

When many amplicons are expected to be amplified simultaneously in the same cell, such as in the case of Fig. 2 (*B. fragilis* CPS operons), it may be advantageous to fuse the P5 sequencing adaptor to each of the locus-specific primers **(Supplementary Table 9)**. In this way, each target amplicon is not competing for the same pool of the P5-adaptor forward primer. The amplicon yield generated for each target will be determined by the amount of target specific primer (containing the P5 adaptor sequence) in the reaction. The downside of this approach is the necessity of synthesizing the long primers (containing the full sequencing adaptor sequence in addition to the target specific sequence) for DoTA-seq.

**Supplementary Note 6:** Recipe for Bacteroides Minimal Media

For 100 mL Bacteroides minimal media (BMM)

| | | | |
|---|---|---|---|
| 1.5 | g | Agar (BD) | (for 1.5% agar plates) |
| 0.5 | g | Glucose (Thermofisher) | |
| 200 | µL | Hemin (2.5 mg/mL) (Sigma) | dissolved in 50 mM NaOH |
| 2 | mg | L-Methionine (Dot Scientific) | |
| 5 | mL | Mineral 3B solution (See below) | |
| 100 | mg | L-cysteine (Dot Scientific) | |
| 150 | µL | FeSO4 (2.8 mg/mL, FeSo4.7H2O) (Alfa Aesar) | add HCl to dissolve |
| Adjust to PH 7.1 | | autoclave | |
| Add | | | |
| 2 | mL | 10% (w/v) Sodium Bicarbonate (Sigma) (Sterilized) | Needed! pH is 3 without this step |

For 100 ml Mineral 3B solution

| | | |
|---|---|---|
| 1.8 | G | KH2PO4 (Alfa Aesar) |
| 1.8 | g | NaCl (Sigma) |
| 40 | mg | MgCl2.6H2O (Sigma) |
| 39 | mg | CaCl2 (Sigma) |
| 2 | mg | CoCl2.6H2O (Sigma) |
| 20 | mg | MnCl2.4H2O (Alfa Aesar) |
| 1 | g | NH4Cl (Fisher) |
| 500 | mg | Na2SO4 (Sigma) |

**Supplementary Note 7:** Recipe for YBHI Media

For 200mL:

7.4 g of BHI Broth (Acumedia Lab)
1 g of Yeast Extract (BD Bacto)
100 mg of D-Cellobiose (Chem-Impex)
200 mg of D-Maltose Monohydrate (Sigma)
100 mg of L-Cysteine (Sigma)
Add 200 mL of MilliQ Water

Autoclave

For plates, add 1.5% w/v Agar
